# Efficient PCA denoising of spatially correlated MRI data

**DOI:** 10.1101/2023.03.29.534707

**Authors:** Rafael Neto Henriques, Andrada Ianuş, Lisa Novello, Jorge Jovicich, Sune N Jespersen, Noam Shemesh

## Abstract

Marčenko-Pastur (MP) PCA denoising is emerging as an effective means for noise suppression in MRI acquisitions with redundant dimensions. However, MP-PCA performance is severely compromised by spatially correlated noise – an issue typically affecting most modern MRI acquisitions – almost to the point of returning the original images with little or no noise removal. In this study, we develop and apply two new strategies that enable efficient and robust denoising even in the presence of severe spatial correlations. This is achieved by measuring a-priori information about the noise variance and combing these estimates with PCA denoising thresholding concepts. The two denoising strategies developed here are: 1) General PCA (GPCA) denoising that uses a-priori noise variance estimates without assuming specific noise distributions; and 2) Threshold PCA (TPCA) denoising which removes noise components with a threshold computed from a-priori estimated noise variance to determine the upper bound of the MP distribution. These strategies were tested in simulations with known ground truth and applied for denoising diffusion MRI data acquired using pre-clinical (16.4T) and clinical (3T) MRI scanners. In synthetic phantoms, MP-PCA failed to denoise spatially correlated data, while GPCA and TPCA correctly classified all signal/noise components. In cases where the noise variance was not accurately estimated (as can be the case in many practical scenarios), TPCA still provides excellent denoising performance. Our experiments in pre-clinical diffusion data with highly corrupted by spatial correlated noise revealed that both GPCA and TPCA robustly denoised the data while MP-PCA denoising failed. In *in vivo* diffusion MRI data acquired on a clinical scanner in healthy subjects, MP-PCA weakly removed noised, while TPCA was found to have the best performance, likely due to misestimations of the noise variance. Thus, our work shows that these novel denoising approaches can strongly benefit future pre-clinical and clinical MRI applications.

## 1. Introduction

The difference in Zeeman energy levels which endows Nuclear Magnetic Resonance (MR) with observable signals is inherently small, typically associated with frequencies in the MHz-GHz range. In MR imaging (MRI), these low energy resonances ensure the safety of the methodology, as strong irradiation is not required to excite spins. However, due to the smallness of the ensuing signal, thermal noise in the receiver coils (Constantinides et al., 1997; Henkelman, 1985) is a significant component of detected data and effectively limits MRI’s spatiotemporal resolution. Furthermore, the potential usefulness of MRI contrasts is often signal-to-noise limited. For example, in diffusion MRI (dMRI) and relaxometry, the already inherently low signals are exacerbated by the need to attenuate the signal for contrast, leading to even more significant losses of spatiotemporal resolution. In functional MRI and methods based on image difference methods (e.g., Magnetic Transfer, Chemical exchange saturation transfer), the contrast to noise is low, and in Magnetic Resonance Spectroscopy (MRS), signals with five orders of magnitude smaller than that of the concentrated water protons are being sought. Thus, suppression of thermal noise effects is of general interest in MRI.

Hardware improvements, such as more efficient coil designs, provide higher signal-to-noise ratio (SNR) from the coil (Kaza et al., 2011; Rodríguez and Medina, 2005; Roemer et al., 1990; Schmitt and Rieger, 2021). More recently, the introduction of cryogenic coils has shown how suppression of thermal noise by a factor of ∼2-10 can produce improvements in image quality (Hall et al., 1991; Kwok and You, 2006; Niendorf et al., 2015; Poirier-Quinot et al., 2008); however, these coils are expensive and difficult to handle (Labbé et al., 2021, 2020). On the other hand, suppression of noise via image processing is a very active field of research, which can potentially provide significant gains in SNR, synergistically with improved coil designs. Several such techniques have been proposed, including standard image-domain smoothing and filtering (Friston et al., 1995; Jones et al., 2005; Jones and Cercignani, 2010), edge-preserving anisotropic filters (Gerig et al., 1992), wavelet transformations (Nowak, 1999; Pižurica et al., 2003), total variation minimization (Knoll et al., 2011), non-local means (Coupé et al., 2008; Manjón et al., 2010, 2008). However, since these techniques rely on assumptions on spatial features of the data, such methods can compromise the image anatomical details by introducing spatial smoothing, blurring, staircase artefacts and other types of image intensity bias (Kay, 2022; Tax et al., 2022; Veraart et al., 2016a, 2016c).

Principal component analysis (PCA) denoising was more recently proposed for a more optimal compromise between noise suppression and preservation of signal information by exploring the MR signal redundancy across multi-dimensional acquisitions (Manjón et al., 2015, 2013; Veraart et al., 2016c). Redundancy exists in many types of MRI methods where larger amount of data samples are acquired relative to the relevant degrees of freedom. This is the case for diffusion MRI data is acquired for e.g., different diffusion gradient directions, different b-values, and/or different diffusion times. In MRI relaxometry, signals are acquired for multiple echoes, and, in functional MRI or functional MRS, redundancy arises from repeating the scans along the paradigm.

In such acquisitions, the denoising is based on a reshaping of data as a *M × N* dimensional matrix, where *M* represents voxels (which can correspond to all image voxels or selected voxels according to a sliding window (Manjón et al., 2013; Veraart et al., 2016c)) and *N*corresponds to the redundant dimension (e.g. along different diffusion gradient acquisitions, echoes, or functional MRI/MRS time points). Following a PCA, a relatively small number of principal components typically carry mainly signal information while the rest carry mainly noise (Manjón et al., 2013) (n.b., the signal components are also perturbed by the noise). In diffusion MRI, for example, it was shown that PCA eigenvalues mainly corresponding to noise can be removed using an empirical threshold calculated based on an a-priori noise variance estimate and a conversion factor that was empirically adjusted according to different acquisition schemes (Manjón et al., 2015, 2013).

Later, the PCA component classification was objectified by Veraart et al. using concepts from random matrix theory (Veraart et al., 2016b, 2016c) and assuming that PCA noise-related eigenvalues are characterized by a Marčenko-Pastur distribution (Marčenko and Pastur, 1967). The Marčenko-Pastur PCA (MP-PCA) denoising has since become one of the most employed algorithms for diffusion MRI pre-processing. The MP-PCA denoising has shown promising results not only for diffusion MRI, e.g., (Ades-Aron et al., 2018; Henriques et al., 2021; Moeller et al., 2021; Olesen et al., 2023; Shemesh, 2018; Tournier et al., 2019; Veraart et al., 2016c), but also for other MRI modalities such as MRI relaxometry (Bazin et al., 2019; Does et al., 2019), functional MRI (Ades-Aron et al., 2020; Adhikari et al., 2019; Diao et al., 2021; Fernandes et al., 2022; Vizioli et al., 2021), MRS (Froeling et al., 2021; Simões et al., 2022), and functional MRS (Mosso et al., 2022). However, it is important to note that MP-PCA denoising schemes assume that noise is spatially uncorrelated – which, as will be shown here, can be violated significantly in real-life data and lead to poor denoising performance.

In typical MRI acquisitions, spatially correlated data can arise from different reconstruction steps such as coil combination and parallel imaging (Aja-Fernández et al., 2015, 2014, 2011; Aja-Fernández and Tristán-Vega, 2012; Constantinides et al., 1997; Pruessmann et al., 1999), k-space gridding, zero-filling and partial Fourier acquisitions (Landman et al., 2009b), spatial smoothing and interpolation during the image reconstruction (Jezzard and Balaban, 1995), and wrapped phase (Moeller et al., 2021; Vizioli et al., 2021), among others. In many cases, data is obtained after multiple pre-processing steps, sometimes irreversibly (e.g., vendor reconstruction), and so spatial noise correlations can be an important confounding factor for MP-PCA denoising.

Here, we explore how spatial correlations impact PCA denoising, and develop strategies that produce much more robust denoising performance even in the presence of spatially correlated data. In particular, we show that a significant improvement in the classification of signal and noise components can be achieved by adding prior information on the noise variance. Two novel PCA denoising strategies are developed: 1) the General PCA (GPCA) denoising uses noise variance estimates determined a-priori without assuming specific noise distribution functions; and 2) the Threshold PCA (TPCA) denoising combines the noise variance prior estimate with MP distribution characteristics to define a threshold for noise component removal. We present the relevant theory, and demonstrate the advantages of these two novel denoising strategies in simulations where ground-truth is known, and diffusion MRI data acquired using a pre-clinical (16.4T) and a clinical (3T) scanner (all code used to produce the figures of this paper is freely available at https://github.com/RafaelNH/PCAdenoising).

## 2. Methods

### 2.1. Theory

#### 2.1.1 PCA Denoising

Let us define a *M × N* matrix ***X*** that contains the *N* redundant measurements (in the case of our dMRI data, these correspond to the different diffusion weighted signals acquired for different gradient directions and b-values) for *M* neighboring voxels typically selected in a sliding window. Each column of ***X*** is subtracted by its mean. Without loss of generality, we consider here matrices with *M* ≥ *N*, but the theory presented below can easily be rewritten for matrices with M < N. The principal component analysis of ***X*** can be performed from the eigen-decomposition of its covariance matrix:

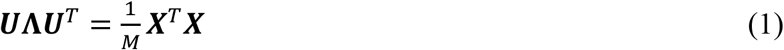

where ***U*** is a *N* × *N* matrix that contains all PCA eigenvectors, and Λ is a *N* × *N* diagonal matrix containing the respective eigenvalues λ. PCA denoising can be achieved by excluding components mainly associated with noise in the eigenvalue spectrum. In pioneering work using PCA to denoise MRI data (Manjón et al., 2015, 2013), this was performed by zeroing eigenvalues lower than a given threshold τ calculated by:

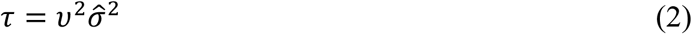

Where 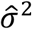 is the noise variance (typically estimated a-priori) and υ is an empirically defined correction factor (Manjón et al., 2015, 2013). Denoised signals for each sliding window are then reconstructed from the eigenvalues (and eigenvectors) surviving the thresholding in PCA space. Denoised datasets can then be fully reconstructed from the middle voxel of each sliding window, or by combining the denoised signals from overlapping voxels from adjacent sliding windows by using, for example, the overcomplete averaging procedure described in (Katkovnik et al., 2009; Manjón et al., 2013). Due to its previous reported advantages (Katkovnik et al., 2009), overcomplete averaging is used for all denoising strategies explored in this study.

Although the pioneering work mentioned above already used a-prior noise variance estimates, latter studies had proposed alternative strategies for component classification to avoid the use of the subjective correction factor (see section 2.1.2). In this study, we show that a-priori noise variance estimates can be used on PCA denoising without employing empirically defined correction factors (see section 2.1.3 and 2.1.4).

#### 2.1.2 MP-PCA Denoising

According to Marčenko and Pastur (1967), a matrix ***X*** populated entirely by pure Gaussian random noise with variance *σ*^2^ in the limit of (*M, N*)_→ ∞_, with *γ* = *N*/*M* fixed, propagates to the PCA eigenvalues λ with the following probability distribution (Marčenko and Pastur, 1967):

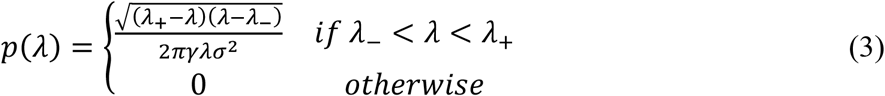

where

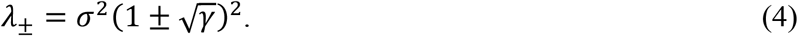

Note that this Marčenko-Pastur (MP) distribution produces non-zero probabilities only between *λ*_−_ and *λ*_+_. From Eq. 4, the width of the MP distribution *λ*_+_ − *λ*_−_ can be related to the noise variance as

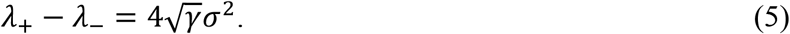

Veraart et al. (Veraart et al., 2016c) proposed to classify eigenvalues mostly related to noise as the maximum number of the smallest eigenvalues λ_*c*_ that best fits the probability distribution in Eq. 3. In practice, this is achieved by a moment-matching method in which larger eigenvalues (carrying significant signal information) are iteratively removed until the mean of the lowest intensity eigenvalues 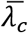 is higher than a noise variance estimated from the MP distribution 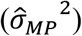, i.e.:

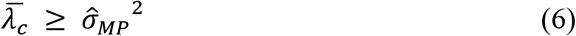

where the noise variance estimate 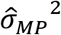 is computed from the MP distribution bandwidth (estimated by the difference between the lower and larger eigenvalues kept in *λ*_*c*_) according to Eq. 5:

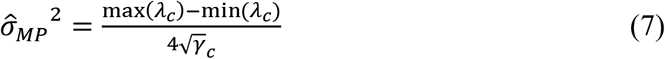

with 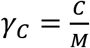 and *C* being the number of eigenvalues classified as being mostly related to noise. Note that MP-PCA denoising does not require prior knowledge about the noise variance since 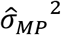 is iteratively calculated by Eq. 7.

#### 2.1.3 General PCA Denoising

The denoising approach described above assumes that the eigenvalue mean 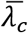 is only larger than 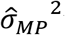 when all components containing relevant signal information are removed from the group of eigenvalues *λ*_*c*_. This assumption holds for independent, uncorrelated entries in the ***X*** matrix. However, this assumption would be violated in typical MRI acquisitions where spatial correlations are introduced by reconstruction in **X**. In that case, as will also be shown below, the original MP-PCA denoising considerably underperforms because it provides an incorrect classification of principal components containing mostly noise and signal information.

Therefore, instead of trying to iteratively estimate the noise variance 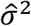 from the Marčenko-Pastur distribution (an estimate that can be corrupted by spatially correlated noise), we introduce here the general PCA denoising approach, which is designed to use an a-priori noise variance estimate 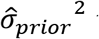 that is obtained independently from the denoising procedure without, however, using empirical conversion factors (Manjón et al., 2015, 2013). In this study, 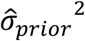 is calculated from MRI data repetitions, but the denoising approaches developed here can be adapted to other 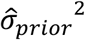 estimation strategies, e.g. (Aja-Fernández et al., 2015; Koay et al., 2009; Landman et al., 2009b, 2009a; Liu et al., 2014; Manjón et al., 2015, 2013; Samsonov and Johnson, 2004; St-Jean et al., 2020) – see, however, the considerations in section 2.1.5.

According to random matrix theory, the mean of PCA eigenvalues is only smaller than the ground-truth noise variance *σ*^2^ when no eigenvalue of significant signal components is improperly classified as noise (Ding et al., 2010; Manjón et al., 2015). Given this, a way to directly use the noise variance prior is to classify eigenvalues of components containing mostly noise by selecting the highest number of smallest eigenvalues whose mean is smaller than 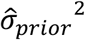, i.e.:

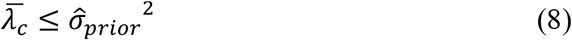

Note that this eigenvalue classification procedure is more general than the criteria used in conventional MP-PCA since it does not rely on specific moments of the Marčenko-Pastur distribution (see supplementary material appendix A) – which is why we refer to this denoising approach as General PCA (GPCA) denoising.

#### 2.1.4 Threshold PCA Denoising

As an alternative to GPCA, the noise variance estimate 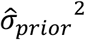 can also be used with an MP distribution-like concept to obtain an objective threshold for PCA component classification. Here, this is achieved by inserting 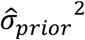 directly into Eq. 4. According to this equation, all noise eigenvalues of a random matrix ***X*** should be lower than *λ*_+_, and thus an eigenvalue threshold criterion for the threshold PCA denoising (TPCA) is here defined as:

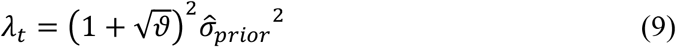

This strategy uses the upper bound *λ*_+_ of the MP distribution, but it does not require the entire distribution to follow exactly the MP itself. In other words the full shape of the eigenvalue probability spectrum is less critical than the upper bound itself. This approach avoids the deleterious effects of spatially correlated noise by avoiding the iterative calculation of two parameters 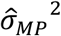 and 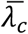 of MP-PCA (c.f. Eq. 6), and rather using the upper bound as the more important metric. Indeed, c.f. Eq. 9, where it is evident that TPCA avoids iterative computation by directly thresholding principal components using the new information from 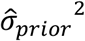which, in a way, accounts for these effects (see considerations below).

#### 2.1.5 Noise Variance Estimation

In modern MRI acquisitions, the noise level may not be uniformly constant across the entire volume (Aja-Fernández et al., 2015; Landman et al., 2009b; Pieciak et al., 2017). Therefore, the noise variance maps in this study are computed at the voxel level. Several techniques have been proposed to calculate these maps from single MRI images, e.g. (Aja-Fernández et al., 2015, 2009; Liu et al., 2014; Pieciak et al., 2017; Tabelow et al., 2015); however, to avoid any assumption about how noise varies spatially across adjacent voxels (Constantinides et al., 1997; Landman et al., 2009b, 2009a; Sodickson et al., 1999), here, noise variance maps were computed from independent images acquired with identical acquisition parameters. For the diffusion MRI datasets analyzed in this study, we used multiple b-value=0 acquisitions and calculated the signal variance across the repetitions (i.e. 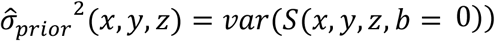). Given the relatively high SNR of b-value = 0 images, this noise estimation strategy is expected to provide accurate noise variance estimates in tissue voxels (Constantinides et al., 1997; Dietrich et al., 2007; Landman et al., 2009b, 2009a; Sodickson et al., 1999); however, in voxels near boundaries (e.g. brain tissues near regions containing cerebrospinal fluid), these noise variance maps may be corrupted (typically overestimated) by image artefacts such as involuntary motion, cardiac pulsation or image intensity drifts (Hansen et al., 2019; Landman et al., 2009b, 2009a; Tournier et al., 2011; Vos et al., 2017). These artefacts can be mitigated in TPCA and GPCA denoising by taking the median of 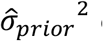 estimates from all voxels of each sliding window instance to achieve an “effective” noise variance estimate, assuming that noise is relatively uniform across each sliding window instance.

#### 2.1.6 Summary of the Main Differences between PCA Denoising Procedures

The three denoising algorithms described above differ in the way they identify PCA components carrying (predominantly) noise and signal. The different algorithms identify the noise by removing the largest eigenvalues until:

1. 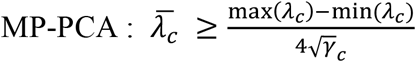
2. 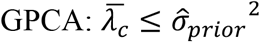
3. 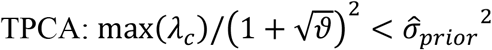

Note that, altogether, the performance of the different algorithms relies on four quantities which can be related to different variance estimates: 1) 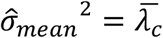 (variance estimated from mean of eigenvalues containing mostly noise); 2) 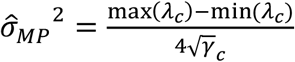 (variance estimates from MP bandwidth); 3) 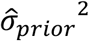 (a-priori variance estimate) and 4) 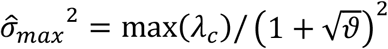 (variance estimate from the maximum eigenvalue containing mostly noise). The effects of spatially correlated noise in these quantities are here explored using simulations (vide infra).

### 2.2. Simulations

In this study, we first show that GPCA and TPCA denoising strategies are more robust to violations of MP-PCA denoising assumptions using simulations where the ground truth number of principal components is known a priori. Two experiments were performed:

#### Experiment 1

A synthetic phantom comprising 12×12 voxels. Signals in this 12×12 grid were generated for 110 different synthetic diffusion-weighted signals, comprising 20 b-value=0 signals and 90 diffusion weighted signals with b-values of 1, 2 and 3 ms/μm^2^ shells with 30 diffusion gradient directions in each shell. This phantom was sub-divided into 9 portions (4×4 voxel regions each) and diffusion-weighted signals were generated using the *Diffusion in Python* (DIPY (Garyfallidis et al., 2014; Henriques et al., 2021)) package according to a forward model containing two well-aligned axially symmetric diffusion tensors. The model parameters of each phantom portion were generated for 9 different principal directions and 9 different volume fractions (evenly sampled from 0.1 to 0.9), while the axial and radial diffusivities (AD and RD, correspondingly) of each component were fixed to AD_1_ = 1.8 μm^2^/ms, AD_2_ = 1.5 μm^2^/ms, RD_1_= 0 μm^2^/ms and RD_2_= 0.5 μm^2^/ms. Diffusion-weighted signals were then corrupted with synthetic Gaussian noise at an SNR level of 30. All voxels were denoised simultaneously (i.e., M=144, N=110 for all algorithms) using the three denoising approaches (MP-PCA (Veraart et al., 2016c), GPCA, TPCA). Since synthetic noise was uniform across voxels and uncorrupted by artifacts, the noise variance for both GPCA and TPCA was set to the voxel averaged 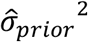 calculated as the variance of b-value=0 signals. To assess the robustness of the denoising towards errors in 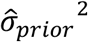, GPCA and TPCA were also run from artificially changing 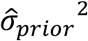 from 50% lower to 400% higher values.

#### Experiment 2

We then assessed the situation in which spatially correlated noise is present. Spatially correlated noise (and signal) was generated by zero-filling 3 columns of the matrices at Fourier space. After corrupting the numerical phantoms with spatially correlated noise, data were denoised using the three denoising approaches. As for the previous experiment, noise variance for both GPCA and TPCA were calculated from the averaged of 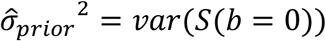 across all image voxels. To assess the generalizability of our results towards different types of spatially correlated noise, simulations with correlated noise due to Gaussian smoothing are presented in supplementary material, appendix B.

Note that, for both experiments, the optimal number of signal components preserved by denoising is known a-priori to be exactly 8 since the phantom was fully constructed from 9 different portions with fixed diffusivities, directions and volume fractions (one signal component must be removed since we subtract the mean of ***X*** before denoising). Moreover, for these simulations, the denoising performance can be directly assessed by comparing signal maps for individual diffusion gradient direction and b-values to their corresponding ground truth maps. The ground truth signals for Experiment 1 corresponded to the synthetic phantom signal before noise corruption, while the ground truth signals for Experiment 2 were computed by zero-filling 3 columns of the original noise free matrices in Fourier space.

### 2.3. Preclinical MRI Experiments

All animal experiments were preapproved by the institutional and national authorities, and carried out according to European Directive 2010/63. A mouse brain (C57BL/6J) was extracted via transcardial perfusion with 4% Paraformaldehyde (PFA), immersed in 4% PFA solution for 24 h, washed in Phosphate-Buffered Saline (PBS) solution for at least 24 h, and then placed on a 10 mm NMR tube filled with Flourinert (Sigma Aldrich, Lisbon, PT), which was sealed using paraffin film.

The MRI experiments were performed on a 16.4 T Bruker Aeon Ascend scanner (Bruker, Karlsruhe, Germany), interfaced with an Avance IIIHD console, and equipped with a gradient system capable of producing up to 3000 mT/m in all directions. A constant temperature of 37°C was maintained throughout the experiments using the probe’s variable temperature capability. Two distinct diffusion-weighted datasets were then acquired using Bruker’s standard “Diffusion Tensor Imaging EPI” sequence for the following diffusion-weighted parameters: 30 gradient directions for b-values 1, 2 and 3 ms/μm^2^ (Δ = 15 ms, δ =1.5 ms), and 20 consecutive b-value=0 acquisitions. Data reconstructed from EPI acquisitions is expected to be corrupted by spatially correlated noise from multiple sources (regridding, partial Fourier factor, multiple shots, including k-space sampling during gradient ramp, etc.). We modulated the amount of spatial correlations by acquiring EPI datasets with parameters optimized to mitigate noise spatial correlations, particularly avoiding k-space undersampling acquisition during EPI’s gradient ramps and without using partial Fourier, which minimize regridding or zero filling. Then, a second dataset was acquired with identical resolution, number of acquisitions, etc., but with large factors inducing spatial correlations, including k-space sampling during gradient ramps (default Bruker’s procedure for acquisition speed) and with a significant phase partial Fourier factor of 6/8. All other acquisition parameters were kept constant: TR/TE = 3000/50 ms, 9 coronal slices, Field of View = 12×12 mm^2^, matrix size 80×80, in-plane voxel resolution of 150×150 μm^2^, slice thickness = 0.7 mm, number of averages = 2, number of segments = 1, double sampling acquisition.

Spatial drifts in the image domain were first corrected using a sub-pixel registration technique (Guizar-Sicairos et al., 2008). Due to the anisotropic voxel sizes (slice thickness > in-plane resolution), MP-PCA, GPCA and TPCA were then applied for each coronal slice separately using a 2D sliding window with 11×11 voxel (M=121, N=110). For all denoising approaches, signals from overlapping windows were combined using the overcomplete averaging procedure described in (Katkovnik et al., 2009; Manjón et al., 2013). Noise variance maps for both GPCA and TPCA were computed from all 20 consecutive b-value=0 acquisitions – final 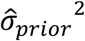 values for each sliding window iteration were taken as the median 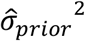 value across all sliding window voxels.

Since diffusion MRI is typically used to compute parametric maps of diffusion properties, in addition to the assessment of the denoising performance of individual diffusion-weighted signals, we also computed standard Diffusional Kurtosis imaging (DKI) (Jensen et al., 2005; Tabesh et al., 2011) metrics reconstructed from both raw and denoised data using the DKI modules implemented in the open-source package *Diffusion in Python* (DIPY) (Henriques et al., 2021).

### 2.4. MRI Experiments Using a Clinical Scanner

Experiments were approved by the Ethical Committee of the University of Trento and the participant signed an informed consent. MRI data was a acquired for a healthy control (male, 54 years) using a 3T MAGNETOM PRISMA scanner (Siemens Healthcare, Erlangen, Germany) equipped with a 64-channel head-neck RF receive coil. Diffusion MRI data was acquired using a monopolar single diffusion encoding EPI PGSE (Feinberg et al., 2010; Moeller et al., 2010; Xu et al., 2013) along 30 diffusion gradient directions for five non-zero b-values = 1, 2, 3, 4.5 and 6 ms/μm2 (Δ = 39.1 ms, δ = 26.3 ms) and 17 interspersed b-value=0 acquisitions. Other acquisition parameters were the following: TR/TE = 4000/80 ms, 63 axial slices, Field of View = 220×220 mm2, matrix size 110×110, isotropic resolution of 2 mm, 6/8 phase partial Fourier, parallel imaging with GRAPPA 2, simultaneous multi-slice factor 3.

MP-PCA, GPCA and TPCA denoising were applied using a 3D sliding window with 9×9×3 voxels (M=243, N=167). To minimize the effects of image artefacts on interspersed b-value=0 acquisitions, initial 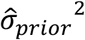 estimates for both GPCA and TPCA were computed from the first five b-value=0 images, and final 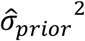 estimates were taken as the median 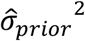 value for each sliding window.

As for the pre-clinical data, denoising performance was assessed on individual diffusion-weighted signals and on diffusion parametric maps. Since DKI fails to represent the diffusion signal decays at high b-values (Chuhutin et al., 2017; Jensen and Helpern, 2010), its maps are reconstructed from the raw/denoised signal for b-value ≤ 3 ms/μm^2^. Additionally, to inspect the preservation of the diffusion MRI angular information at high b-value data, the data for b-values = 4.5 and 6 ms/μm^2^ and respective interspersed b-value=0 acquisitions were subsampled and used to reconstruct Q-ball diffusion orientation distribution functions (dODFs) using the constant solid angle procedure (Aganj et al., 2010) also implemented in DIPY.

## 3. Results

### 3.1. Numerical Simulations

#### 3.1.1 Scenario A: Spatially Uncorrelated Noise

To provide ground truth and validate that all methods perform similarly under ideal conditions, simulations for a phantom with spatially uncorrelated noise are shown in Fig. 1. Ground truth, raw noisy and denoised diffusion-weighted signals for the first diffusion gradient direction of the highest diffusion gradient magnitude simulated (b = 3 ms/μm^2^) are shown in upper panels (Fig. 1A-E). In Fig. 1F (and Fig. 1G for zoomed plotted), the quantities 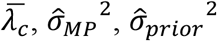 and 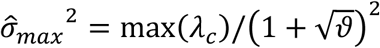 evaluated by the denoising algorithms to classify PCA eigenvalues are plotted. In Fig. 1E and Fig. 1G, the number of noise components classified by the denoising algorithms are marked by the vertical lines (cyan, green, and orange vertical lines for MP-PCA, GPCA, and TPCA, respectively). All algorithms successfully classified the 102 components containing mostly noise and the 8 signal components when noise was spatially uncorrelated as evident by the coincidence of the vertical lines (c.f. black arrows in Fig. 1A3). Fig. 1H shows the eigenvalue spectrum reconstructed by repeating the simulations 1000 times and the theoretical MP distribution with equal eigenvalue variance (plotted in log scale for better visalization of the lower probabilities for higher eigenvalues). As expected, the reconstructed spectrum matches the theoretical MP distribution when noise is spatially uncorrelated. For all three PCA denoising algorithms, the threshold average across the 1000 simulation repetitions approaches the distribution upper bound (vertical lines in Fig. 1H). Since all three algorithms classified identical number of signal components, their performances are qualitatively indistinguishable from each other (c.f. Fig. 1A-E) – the same can be observed for DKI maps reconstructed from the denoised phantoms (Fig. 3A-B). To test how these methods perform in case of 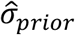 overestimation, the number of components classified as mostly containing noise are plotted in Fig. 1I for different 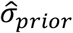 underestimation and overestimation factors. TPCA is more robust to 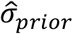 misestimation than GPCA.

**Fig. 1.**
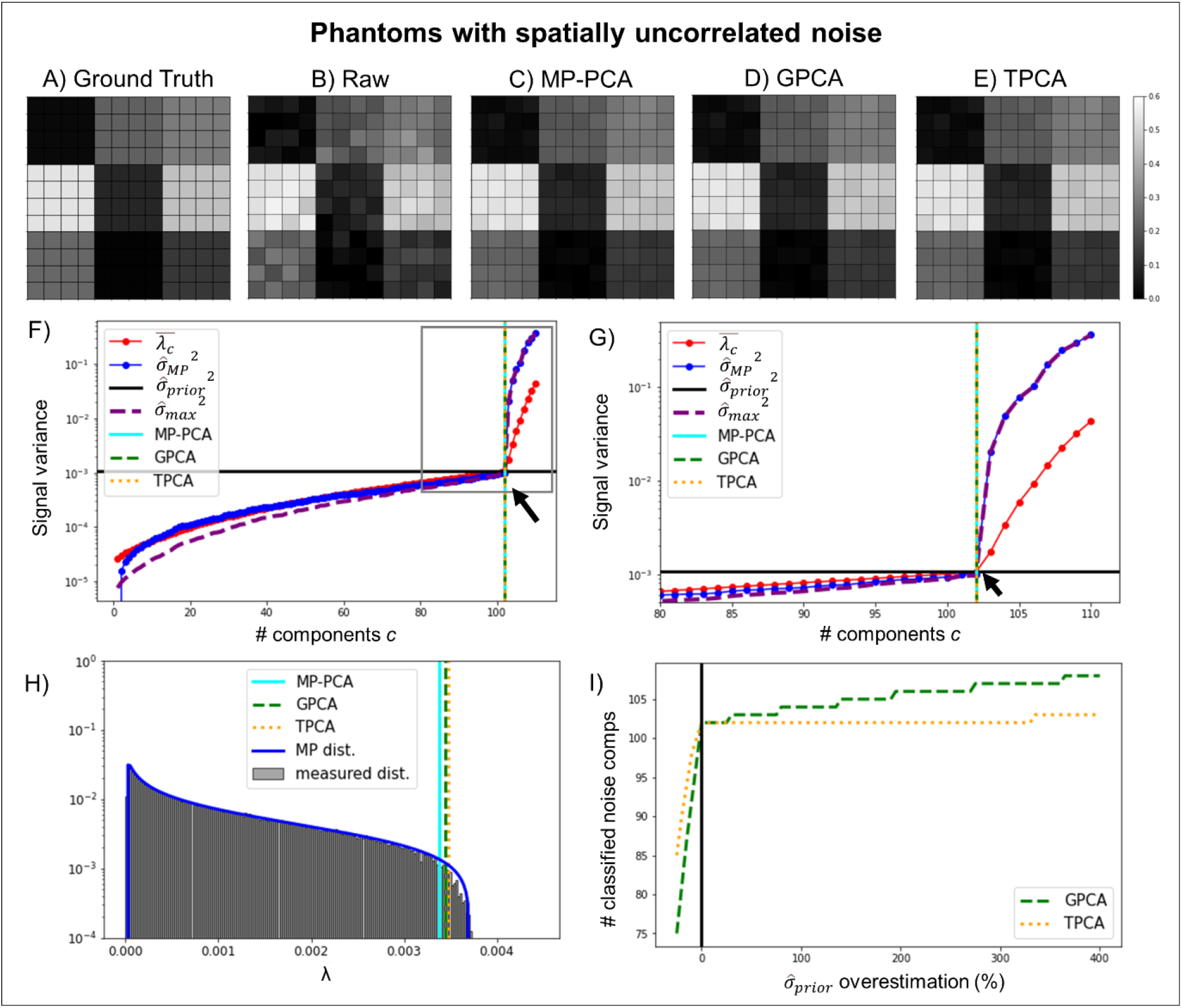
Simulations of denoising performance in a phantom with uncorrelated noise. Representative ground truth, noise free **(A)**, and noise corrupted signals **(B)**, for the first diffusion gradient direction of the highest diffusion gradient intensity are shown in panels and respectively, while denoised signals for the MP-PCA **(C)**, GPCA **(D)**, and TPCA **(E)** denoising algorithms. **(F)** Parameters assessed by the denoising algorithms plotted as a function of the number of lower eigenvalues potentially considered as noise in panel - thresholds for the MP-PCA, GPCA and TPCA are plotted by the cyan solid, green dashed, and orange vertical lines respectively (black arrow point to the ground truth number of signal components, i.e., 102). **(G)** Zoomed plot of the parameters assessed by the denoising algorithms. **(H)** Reconstructed eigenvalue spectrum for 1000 trials and respective theoretical MP distribution for identical eigenvalue variances are shown in panel– the mean thresholds for the MP-PCA, GPCA and TPCA computed as the threshold average across the 1000 repetitions are plotted by the cyan solid, green dashed, and orange vertical lines respectively. **(I)** The number of classified noise components for both GPCA and TPCA as a function of the percentage overestimation of noise standard deviation. All algorithms produced identical denoising performances when noise is spatially uncorrelated.

#### 3.1.2 Scenario B: Spatially Correlated Noise

Next, we investigate the phantom with spatial correlations (Fig. 2) by zero-filling 3 data columns in Fourier space, which interpolates the data in image domain and creates strong spatial correlations (equivalent results for another scenario creating spatial correlations, namely due to smoothing in image domain, are shown in supplementary Fig. S1). Ground truth, raw noisy, and denoised diffusion-weighted signals for the first diffusion gradient direction of the highest diffusion gradient magnitude simulated (b= 3ms/μm^2^) are presented in Fig. 2A-E. The MP-PCA denoising classification criterion 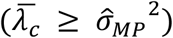 identified only a small number of PCA eigenvalues as mostly carrying noise (solid cyan line in Fig. 2E). That is, the classification procedure underestimated the number of noise components, thereby negatively impacting denoising performance.

**Fig. 2.**
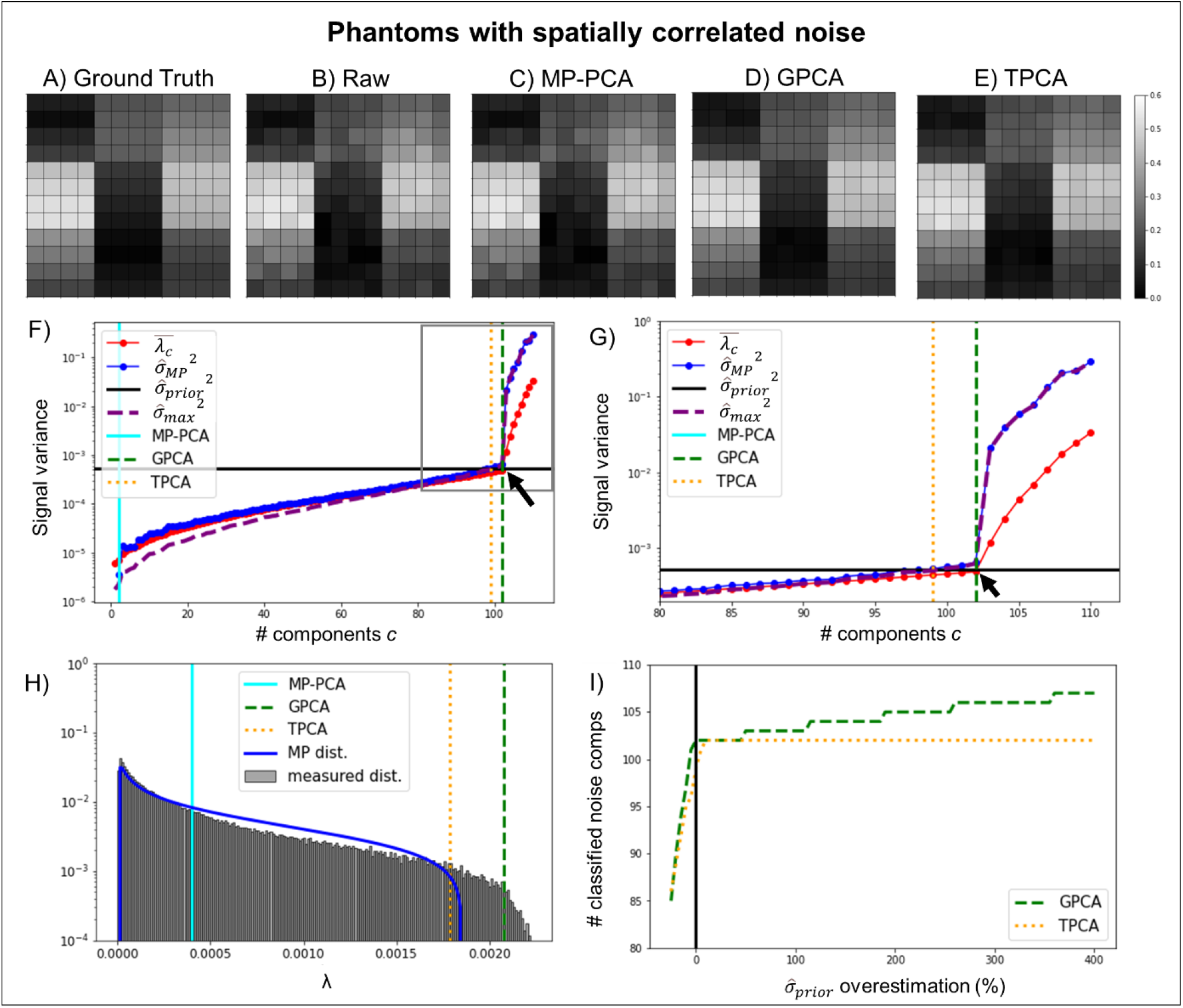
Simulations of denoising performance in a phantom with correlated noise. Correlations were induced by zero filling in k-space. Representative ground truth noise free **(A)** and noise corrupted **(B)** signals for the first diffusion gradient direction of the highest diffusion gradient intensity alongside denoised signals for the MP-PCA **(C)**, GPCA **(D)** and TPCA **(E). (F)** Parameters assessed by the denoising algorithms are plotted as a function of the number of lower eigenvalues potentially considered as noise. Thresholds for the MP-PCA, GPCA and TPCA are plotted by the cyan solid, green dashed, and orange vertical lines respectively (black arrow point to the ground truth number of signal components, i.e., 102). **(G)** Zoomed plot of the parameters assessed by the denoising algorithms. **(H)** Reconstructed eigenvalue spectrum for 1000 instantiations and respective theoretical MP distribution for identical eigenvalue variances. The mean thresholds for the MP-PCA, GPCA and TPCA computed as the threshold average across the 1000 repetitions are plotted by the cyan solid, green dashed, and orange vertical lines respectively. **(I)** The number of classified noise components for both GPCA and TPCA as a function of the percentage overestimation of noise standard deviation. Thus, GPCA and TPCA denoising algorithms outperform MP-PCA denoising when noise is spatially correlated.

In contrast to the conventional MP-PCA, GPCA denoising correctly classified all 102 components containing mostly noise and 8 signal components (dashed orange line in Fig. 2E and Fig. 2G) based on the criterion 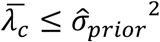. TPCA misclassified 2 ground truth noise components as being significant signal components (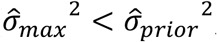, dashed green line in Fig. 2E and Fig. 2G), but still exhibited good denoising performance without removing signal components. Fig. 2H shows that the bandwidth of the eigenvalue spectrum for eigenvalues of spatially correlated noise is increased relative to the theoretical MP distribution with equal eigenvalue variance. Qualitatively, while the denoised signals from MP-PCA (Fig. 2C) were nearly identical to the raw non-denoised signals (Fig. 2B), the denoised signals from both GPCA (Fig. 2D) and TPCA (Fig. 2E) corresponded much better to their respective ground truth maps (Fig. 2A). Similar observations were made for the extracted DKI parameters (Fig. 3C-D). As for the previous scenario, TPCA is more robust to 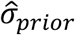 misestimation than GPCA, even for spatially correlated noise (Fig. 2I).

**Fig. 3.**
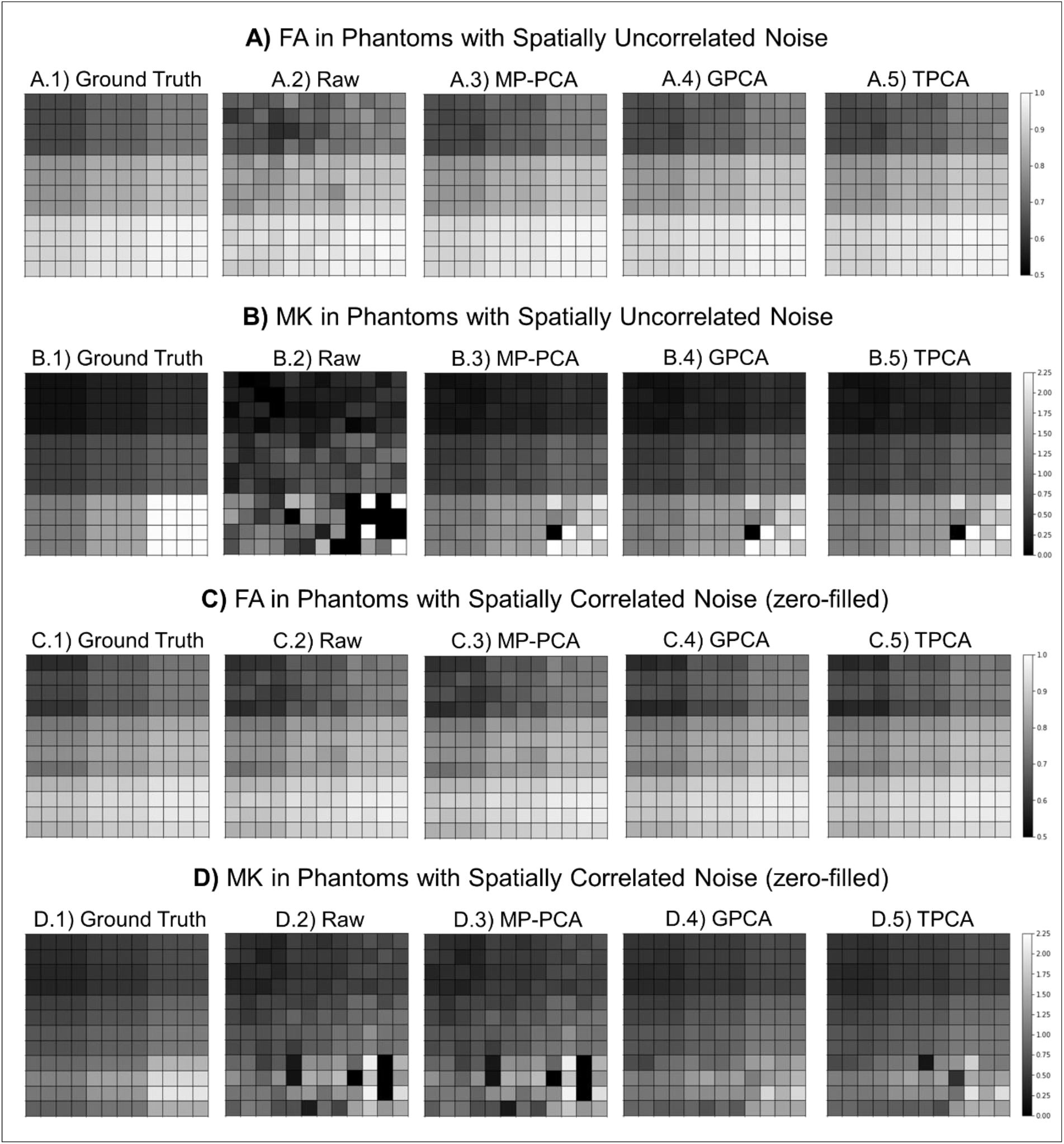
Simulated denoising performance in parametric FA and MK estimates. **(A)** FA estimates for simulations with spatially uncorrelated noise; **(B)** MK estimates for simulations with spatially uncorrelated noise; **(C)** MK estimates for simulations with spatially correlated noise; and **(D)** FA estimates in simulations with spatially correlated noise. MK estimates in simulations with spatially correlated noise. From left to right, panels show the maps computed from ground truth signals (A1, B1, C1, D1), the maps computed from noise corrupted signals (A2, B2, C2, D2), the maps for computed from MP-PCA denoised signals (A3, B3, C3, D3), the maps for computed from GPCA denoised signals (A4, B4, C4, D4), and the maps for computed from TPCA denoised signals (A5, B5, C5, D5). GPCA provides the best denoising performance in this case where the noise variance is accurately estimated.

### 3.2. Preclinical MRI Experiments

#### 3.2.1 Data Acquired with Small Spatial Correlations

Results for the diffusion-weighted data acquired with parameters adjusted to minimize noise spatial correlations are presented in Fig. 4A. From left to right, the upper panel (Fig. 4A1) shows a raw representative image acquired with a b-value of 3 ms/μm^2^, the initial noise standard deviation 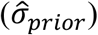 map before performing the median filtering across the voxels of the PCA denoising sliding windows, and the final noise standard deviation 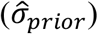 map after selecting the median value of each sliding window, which was used in the subsequent denoising procedures. The initial noise standard deviation map (middle image of Fig. 4A1) was spatially uniform across the brain, except for higher values near brain edges likely due to small image intensity drifts and residual spatial drifts (yellow arrows in Fig. 4A1). These higher noise variance values were successfully attenuated after selecting the median values across sliding windows (right image of Fig. 4A1). The lower panel (Fig. 4A2) shows the corresponding denoised diffusion-weighted image, the denoising residuals computed as the subtraction between the denoised and raw representative image, and number of classified signal components for all three denoising algorithms tested – MP-PCA, GPCA and TPCA from top to bottom. For this dataset with few spatial correlations, all denoising algorithms performed similarly. For instance, all strategies uniformly classified around 10 PCA signal components across the brain (Fig. 4A2 and Fig. 4A3), except in regions near the brain ventricles and boundaries in which the MP-PCA classified a larger number of signal components (Fig. 4A2). Still, the residual maps are clearly similar and show no structure (Fig. 4A2).

**Fig. 4.**
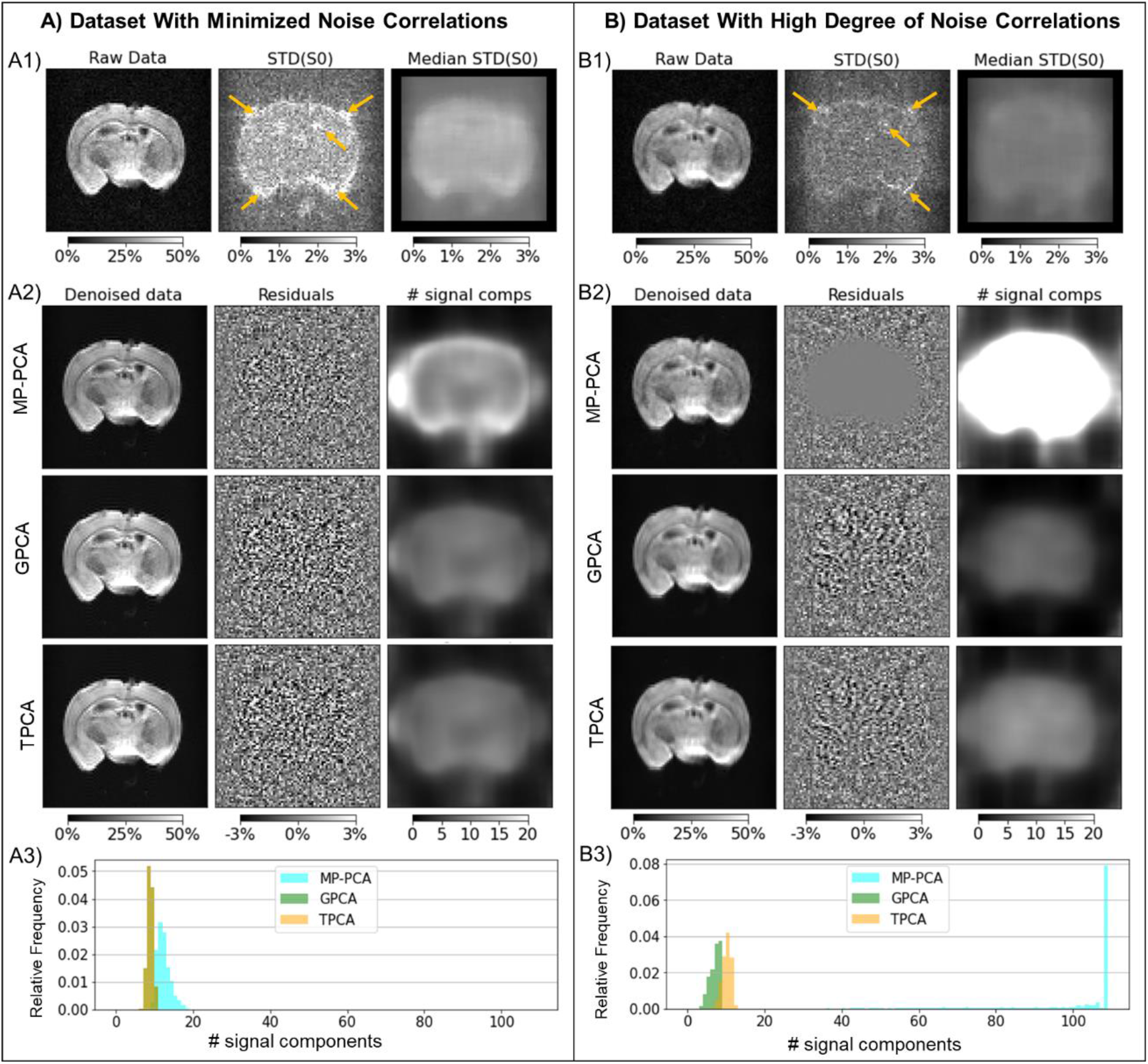
Denoising performance in preclinical datasets. **(A)** Uncorrelated noise scenario and **(B)** acquisition with much higher noise correlations. From left to right, the upper panels **(A1/B1)** show a representative diffusion MRI slice for a selected gradient direction acquired with b-value = 3ms/μm^2^, the initial noise standard deviation (STD) maps computed as the standard deviation of the signals across the data for different b-value=0 acquisitions, and the finally noise STD maps computed after selecting the median values across the voxels of the PCA sliding windows (yellow arrows points regions of high STD noise estimates). From left to right, the lower panels **(A2/B2)** show the denoised data for the selected diffusion MRI image, the denoising residuals computed as the difference between denoised and raw data, and the number of PCA signal components preserved by each denoising algorithm (upper to lower panels shows the results for MP-PCA, GPCA, and TPCA respectively). **(A3/B3)** shows the relative frequencies of the number of PCA signal components preserved by each denoising algorithm across all brain voxels (note in A3 GPCA relative frequencies are overlapped by TPCA frequencies). The map intensities for raw and denoised data as well as noise STD estimates and denoising residuals are displayed in reference to the mean b=0 signals across all brain voxels (a value of 100% corresponds to values equal to that reference value). All denoising algorithms have similar performances on data with minimal spatially correlated noise **(A)**; however, MP-PCA fails to denoise data highly corrupted by spatially correlated noise while both GPCA and TPCA maintain their performance **(B)**.

#### 3.2.2 Dataset Acquired with Large Noise Correlations

Fig. 4B shows the results for diffusion-weighted data acquired with factors inducing significant spatial correlations, including partial Fourier and signal acquisition during EPI’s gradient ramp (no ramp compensation). These are much more commonly used settings than those shown in Fig. 4A. Representative images are shown in Fig. 4B1 left, while noise standard deviation maps 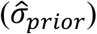 before and after selecting the median values across the voxels of PCA denoising sliding windows are shown in the middle and right images of Fig. 4B1.

In this type of data with spatial noise correlations, the MP-PCA denoising failed: residuals are close to zero across all brain voxels and more than 40 components were classified as significant signal components (upper middle and right images of Fig. 4B2 and Fig. 4B3). We then tested whether GPCA and TPCA could outperform the MP-PCA. Indeed, a uniform denoising performance across the brain and background regions was clearly observed for these two algorithms (lower rows of panels in Fig. 4B2). As the previous dataset in Fig. 4A, GPCA and TPCA preserve around 10 PCA components across the brain (Fig. 4B2 right panels and Fig. 4B3).

#### 3.2.3 DKI Parametric Maps

Fig. 5 shows the parametric maps for four DKI metrics: A) Fractional Anisotropy (FA); B) Mean Kurtosis (MK); C) Radial Kurtosis (RK); and D) Axial Kurtosis (AK). These were reconstructed from the data acquired with factors inducing spatial correlations (for the counterpart with much less correlations, c.f. supplementary Fig. S2). Other diffusion parameters are also given in supplementary Fig. S3. For each panel in Fig. 5, DKI maps were separately computed from the raw data and data denoised by MP-PCA, GPCA, and TPCA. For a better inspection of the denoising performance, these panels display maps for an entire representative data slice (upper panels A1, B1, C1, and D1) and for the zoomed area marked by the red box (lower panels A2, B2, C2, and D2). Since MP-PCA failed to effectively suppress noise for this dataset (c.f. Fig. 4B), DKI maps appear very similar between the non-denoised and MP-PCA denoised data. By contrast, DKI maps computed from data denoised by the GPCA and TPCA algorithms evidence better maps, decreases in spurious fluctuations (orange arrows in Fig. 5A-D), and preservation of contrast between white and grey matter tissues (cyan arrows in Fig. 5A-D). In addition, qualitatively, GPCA and TPCA evidenced very similar denoising performances.

**Fig. 5.**
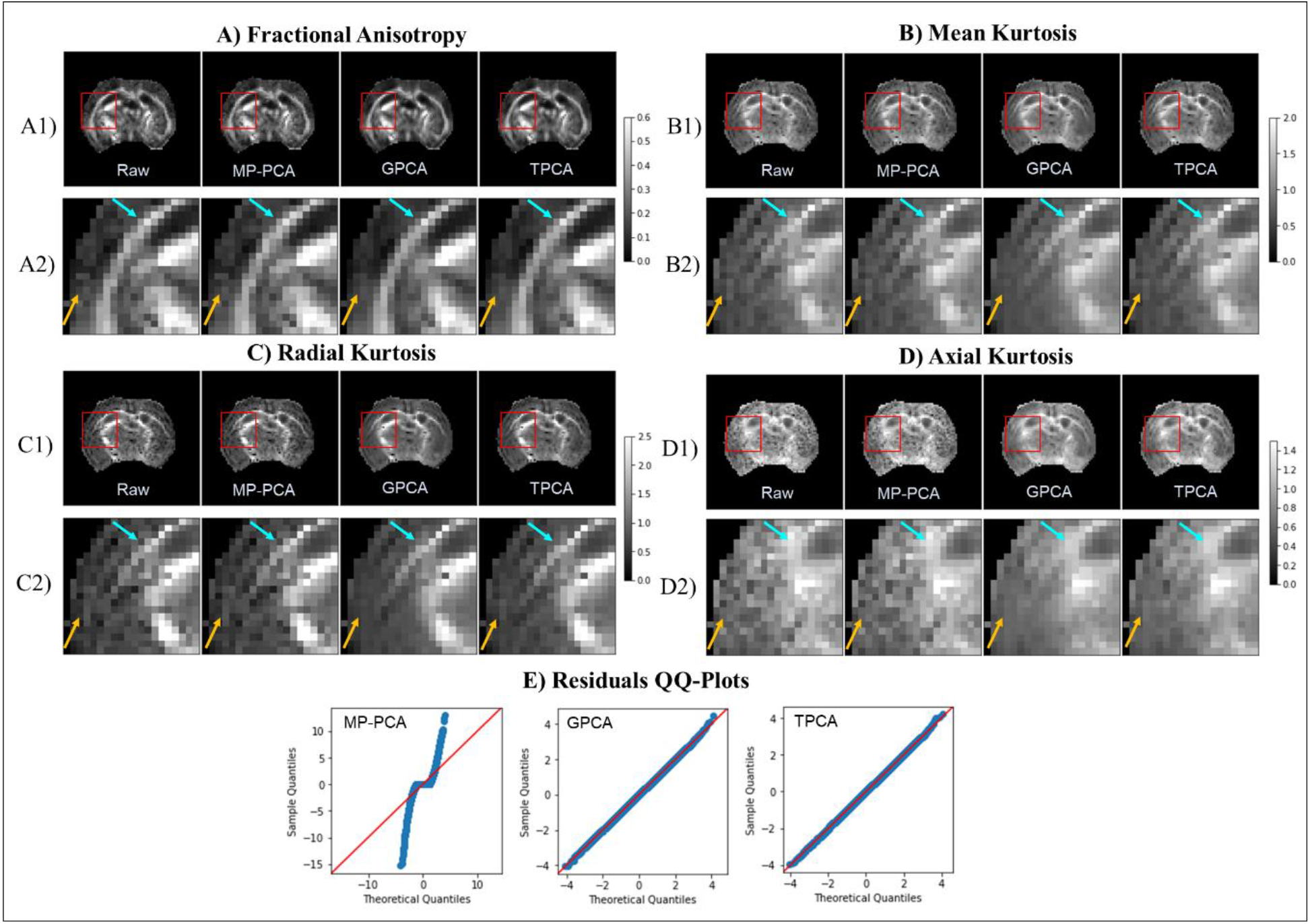
DKI maps for the pre-clinical dataset highly corrupted by spatially correlated noise and QQ-plots of the denoising residuals: **(A)** Fractional Anisotropy; **(B)** Mean Kurtosis; **(C)** Radial Kurtosis; **(D)** Axial Kurtosis; **(E)** QQ-plots of the denoising residuals. For each DKI quantity, images are displayed for an entire representative axial slice (A1, B1, C1, D1) and for the zoomed area marked by the red box (A2, B2, C2, D2). From left to right, DKI maps are displayed for the raw, MP-PCA denoised, GPCA denoised and TPCA denoised data (orange arrows point to areas where noise estimate fluctuations are visually reduced, while blue arrows points to boundary areas between grey and white matter). QQ-plots of the denoising residuals for the selected zoomed region are show panel E. Note that, even when noise is highly spatially correlated, reconstruction DKI are improved on data denoised by the GPCA and TPCA procedures.

To better quantify these effects, QQ-plots (Wilk and Gnanadesikan, 1968) of the denoising residuals for the selected zoomed region of interest are shown in Fig. 4E. For both GPCA and TPCA, the sampled residual quantiles closely matched the theoretical quantiles of a Gaussian distribution, as evident from the linear nature of the plot. By contrast, substantial deviations between MP-PCA sampled residual quantile and the theoretical Gaussian distribution reference reflect the poor denoising performance of MP-PCA for the pre-clinical dataset acquired with no concerns on spatially correlated noise.

### 3.3. MRI Experiments Using a Clinical Scanner

#### 3.3.1 Diffusion-weighted Images

Fig. 6 shows results from a healthy volunteer. Fig. 6A shows the mean signal of the 5 repeating b =0 acquisitions (Fig. 6A1), the initial 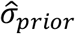 map computed from these b = 0 acquisitions (Fig. 6A2), and the final “effective 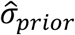” map after selecting the median values from PCA denoising sliding windows (Fig. 6A3). Higher initial 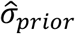 estimates are observed in regions near the ventricles and cortical boundaries (e.g., yellow arrows in Fig. 6A2), likely an effect of signal fluctuations in these regions due to pulsation artefacts (Tournier et al., 2011). These higher noise variance estimates are suppressed but not completely removed upon median filtered 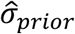 maps (e.g., yellow arrows in Fig. 6A3).

**Fig. 6.**
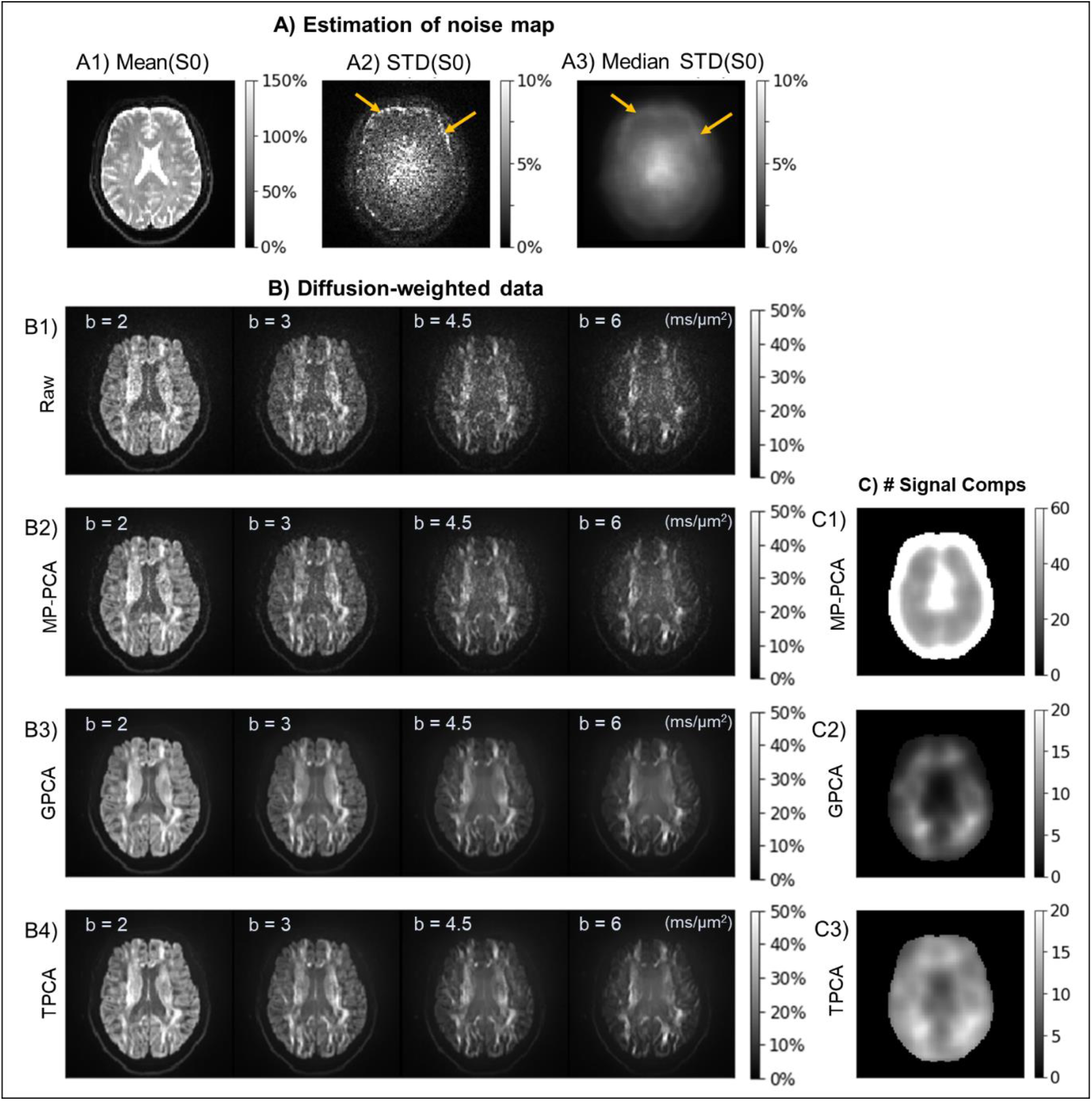
Denoising performance in diffusion-weighted data acquired using a clinical scanner. **(A)** Images related to the noise prior estimation: representative slice of the averaged signals of the 5 first b-value=0 acquisitions (A1); initial noise standard deviation (STD) maps computed as the standard deviation of the 5 first b-value=0 acquisitions (A2); and the final noise STD maps computed after selecting the median values across the voxels of the PCA sliding windows (A3) - yellow arrows points regions of high STD noise estimates. (**B)** Representative slice of the diffusion-weighted data for a gradient direction near **v** = [1, 0. 0] and for b-value = 2, 3, 4.5, and 6 ms/μm^2^ (from left to right) before denoising (B1) and after denoising using the MP-PCA (B2), GPCA (B3), and TPCA (B4) procedures. **(C)** Number of PCA signal components preserved by the three denoising algorithms: MP-PCA (C1); GPCA (C2); and TPCA (C3). The map intensities for raw and denoised data as well as noise STD estimates and denoising residuals are displayed in reference to the mean b-value=0 signals across all brain voxels (a value of 100% corresponds to values equal to that reference value). GPCA and TPCA have superior denoising performance over MP-PCA in a typical high b-value diffusion-weighted dataset.

Representative images of the raw and denoised diffusion-weighted data for the 4 higher non-zero b-values (b-value = 2, 3, 4.5, and 6 ms/μm^2^) are shown in Fig. 6B. The number of preserved signal component maps for all three denoising procedures is shown in Fig. 6C. For this dataset, GPCA and TPCA improved denoising performance over the MP-PCA (Fig. 6B2 vs Fig. 6B3/Fig. 6B4). Specifically, while MP-PCA classified over 40 of the 167 PCA components as significant signal components (Fig. 6C1), GPCA and TPCA classified around 10 to 20 signal components across all brain regions (Fig. 6C2, 6C3). TPCA tended to classify slightly more signal components than GPCA. Denoising residuals for each procedure are shown in Supplementary Fig. S4. GPCA produced higher amplitude residuals. Qualitatively, no structural information is observed in these residual maps.

#### 3.3.2 Diffusion Parametric Maps

Fig. 7 shows sample DKI parametric maps computed from raw and denoised human brain data. As for the pre-clinical data presented above, better noise suppression was observed in diffusional kurtosis quantities (i.e., MK, RK, AK) for data denoised by GPCA and TPCA procedures compared with the MP-PCA counterpart. Note for example the reduction of implausible negative kurtosis values in MK and RK maps for GPCA and TPCA denoised data (Fig. 7B and Fig. 7C). After a closer inspection of DKI estimates in regions near brain edges (regions of interest marked by the yellow boxes), decreases in FA and RK were observed for the data denoised by the GPCA denoising strategy (red arrows in Fig. 7E and Fig. 7F), suggesting that some true signal was also removed. For both MP-PCA and GPCA denoising, QQ-plots of the denoising residuals in this region of interest show substantial deviations between the sampled residual quantiles and the theoretical Gaussian distribution reference (Fig. 7G). While sampled residual quantile deviations for MP-PCA reflect poor removal of components mostly containing noise, deviations for GPCA indicate loss of signal information. The closer match between sampled residual quantiles and the theoretical Gaussian quantiles for TPCA suggests that this algorithm provides a better compromise between noise suppression and signal preservation.

**Fig. 7.**
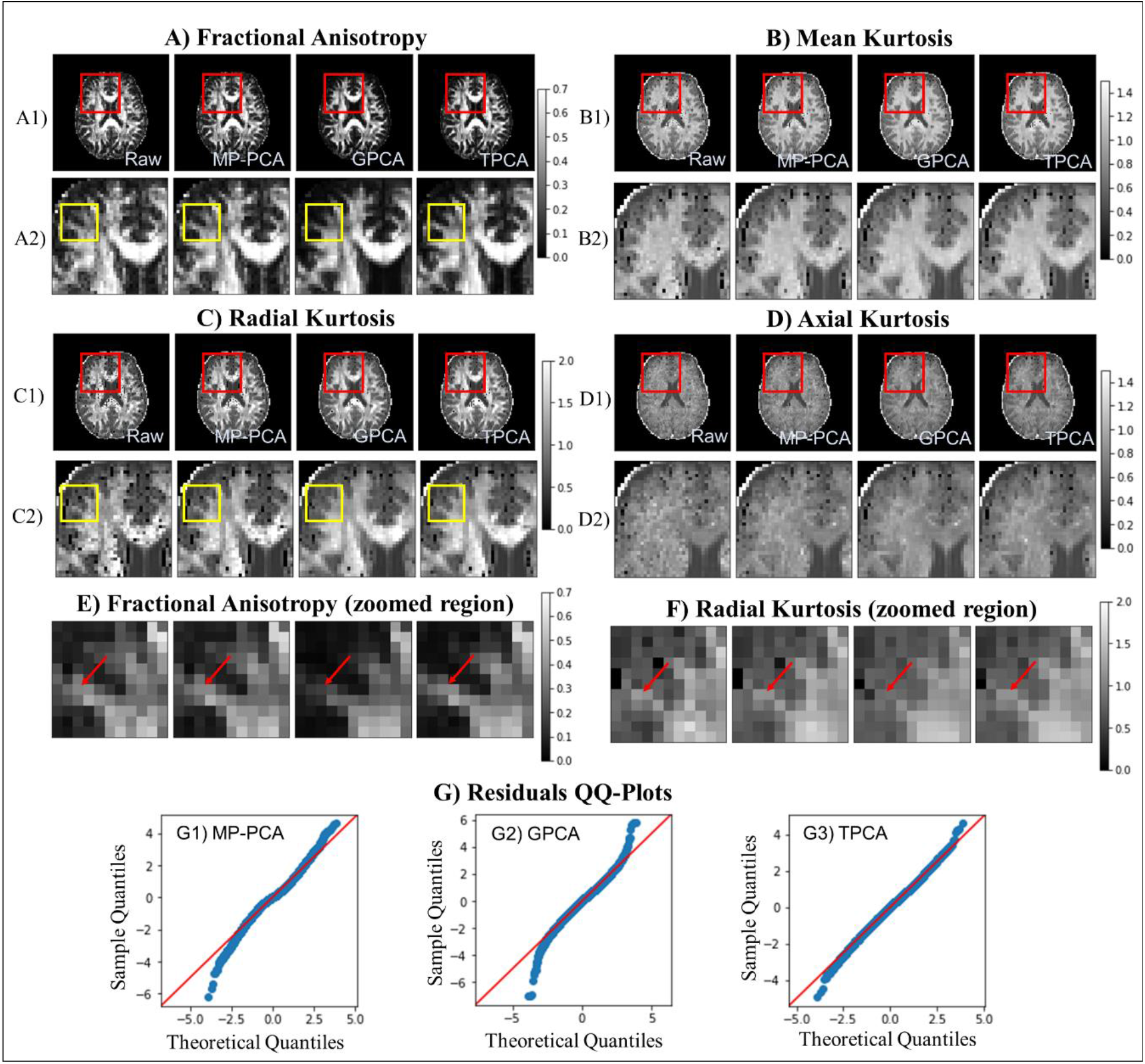
DKI maps for the diffusion-weighted human data computed for all data with b-value ≤ 3 ms/μm^2^ and QQ-plots of the denoising residuals. **(A)** Fractional Anisotropy; **(B)** Mean Kurtosis; **(C)** Radial Kurtosis; **(D)** Axial Kurtosis; **(E)** Fractional Anisotropy in a zoomed problematic region of interest (yellow regions marked in previous maps); **(F)** Zoomed Radial Kurtosis in a zoomed problematic area of interest (yellow regions marked in previous maps); and **(G)** QQ-plots of the denoising residuals. For panels A-D, images are displayed for an entire representative axial slice (A1, B1, C1, D1) and for the zoomed area marked by the red box (A2, B2, C2, D2). From left to right in panels A-F, DKI maps are displayed for the raw, MP-PCA denoised, GPCA denoised and TPCA denoised data. QQ-plots of the denoising residuals for the selected zoomed yellow region are show panel E. Both GPCA and TPCA enhance the quality of DKI reconstruction for the human data. However, while GPCA induces decreased FA and RK estimates for regions near brain edges, TPCA provides an optimal compromise between noise suppression and signal preservation at these problematic regions.

To further examine the potential utility of these denoising schemes, general fractional anisotropies (GFA) maps and diffusion orientation distribution functions (ODF) were computed from Q-ball raw and denoised human brain data acquired at higher b-values (Fig. 8). Consistent with the DKI results presented above, MP-PCA denoised the data quite poorly, while GPCA and TPCA removed noise efficiently (Fig. 8A), but with GPCA apparently removing some signal components (c.f. red arrow in Fig. 8A). To better visualize ODF differences, two regions of interest were manually defined (Fig. 8B): (1) a region comprising voxels where white matter fibers are expected to cross (ROI 1); and (2) regions comprising voxels of lateral corpus callosum fiber projections where the previous FA and RK for GPCA denoised data showed decreased values (ROI 2). In the latter, the ODFs reconstructed from the raw and MP-PCA denoised data show implausible multiple lobes. Consequently, these reconstructions poorly characterized the differences expected between ODFs in crossing regions and the lateral single fiber corpus callosum projections (c.f. first two columns of images in Fig. 8C and Fig. 8D). The ODF reconstructed from the GPCA and TPCA denoised data shows the expected multiple and single lobes for ROI 1 and ROI 2 respectively (c.f. last two columns of images in Fig. 8C and Fig. 8D); ODFs reconstructed from TPCA show, however, shaper profiles, indicating that this denoising procedure better conserves diffusion angular information. GPCA and TPCA exhibited higher reproducibility of ODF profiles across the data from the two independent b-values 4.5 and 6 ms/μm^2^. Note that ODF profiles between b-values 4.5 and 6 ms/μm^2^ data are only consistent when denoised by GPCA and TPCA algorithms (Fig.8C and Fig. 8D), but not with MP-PCA.

**Fig. 8.**
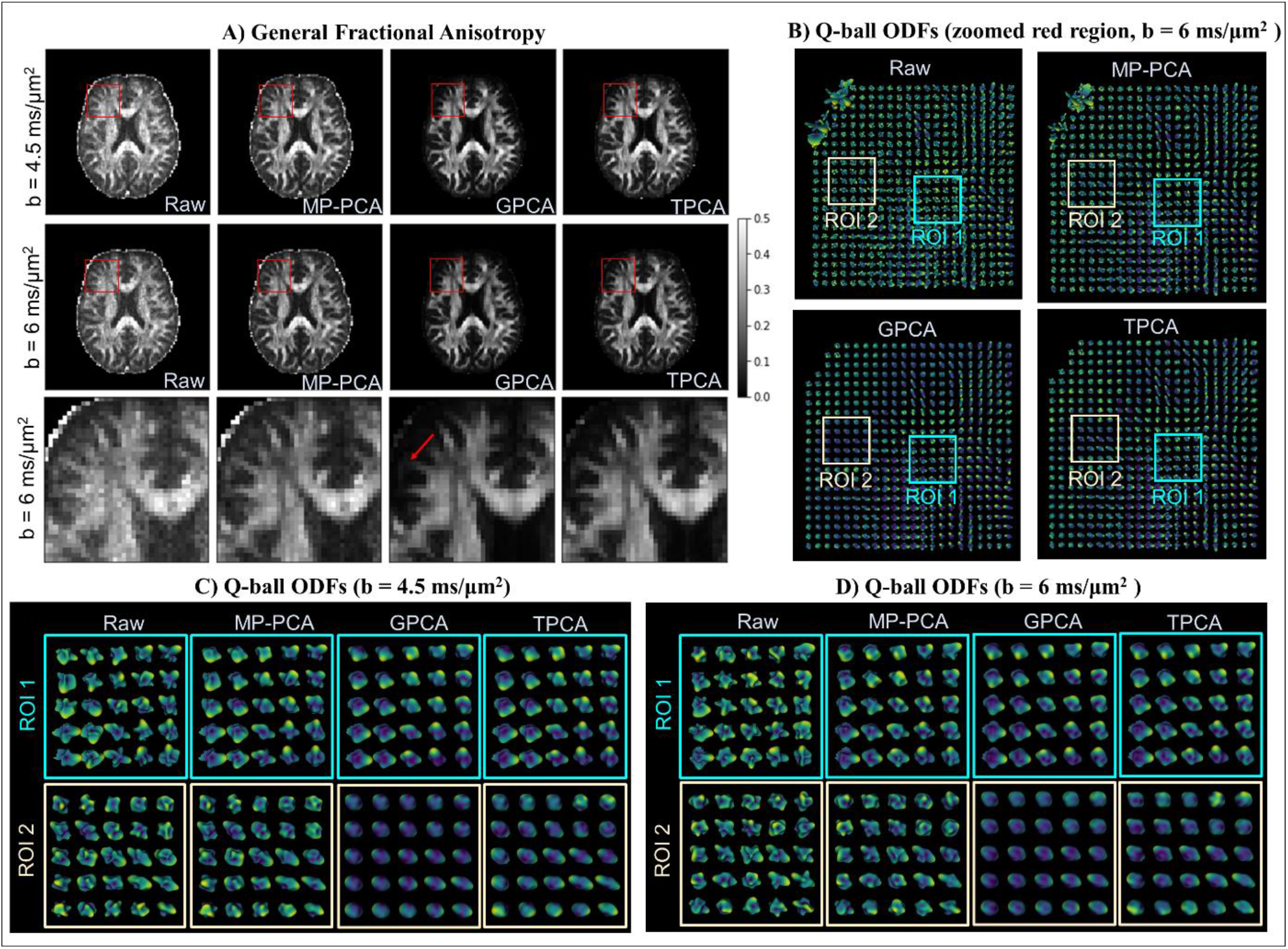
Q-ball GFA maps and ODF reconstruction for the human diffusion-weighted data independently computed for the two higher b-values 4.5 and 6 ms/μm^2^. **(A)** General Fractional Anisotropy maps for the raw and denoised data acquired for b-value=4.5 ms/μm^2^ (upper full slice images) and b-value=6 ms/μm^2^ (lower full slice and zoomed images) data – red arrow indicates a region with artificially decreases diffusion anisotropy. **(B)** ODFs reconstructed for the b-value = 6 ms/μm^2^ data in the zoomed red regions marked on the upper images in panel A. **(C)** ODFs reconstructed for the b-value = 4.5 ms/μm^2^ in the two regions of interest defined in panel B. **(D)** ODFs reconstructed for the b-value = 6 ms/μm^2^ in the two regions of interest defined in panel B. GPCA and TPCA reduce implausible lobes in Q-ball ODF reconstructions. ODF reconstructions from TPCA, however, produces sharper profiles in regions near brain edges (ROI 2).

## 4. Discussion

Denoising has become a critical component of data analysis in quantitative neuroimaging (Ades-Aron et al., 2018; Adhikari et al., 2019; Diao et al., 2021; Kay, 2022; Kay et al., 2013; Tax et al., 2022). The existing approaches vary from subjective (PCA thresholding) (Manjón et al., 2015, 2013) to more objective (MP-PCA) approaches (Ades-Aron et al., 2020; Does et al., 2019; Fernandes et al., 2022; Moeller et al., 2021; Mosso et al., 2022; Olesen et al., 2023; Simões et al., 2022; Veraart et al., 2016b, 2016c; Vizioli et al., 2021). The objectivity of the latter approach is a great advantage, ensuring that only noise is removed while signal components are untouched, thereby leading to data that is denoised but not smoothed or otherwise corrupted. The existing recommendations for the application of MP-PCA denoising already acknowledge the issue of spatial correlations and suggest applying the denoising routing at the earliest step of signal reconstruction and processing. However, in practice, this is seldom possible, whether due to the vendor’s data output format (e.g. coil-combined magnitude images) and/or due to the relative difficulty in executing complex image reconstruction pipeline for many users. However, as shown explicitly in this study, correlations can exist in reconstructed datasets – whether due to commonly used partial Fourier encoding, interpolation, or gridding of data sampled in a non-cartesian way – and, as shown here (and anticipated in previous work), they significantly degrade the performance of MP-PCA. Hence, in this study we sought to develop and explore PCA denoising approaches that increase the robustness of the classification of components containing mostly noise or signal to the effects of spatially correlated noise. Using an additional explicit measurement of the noise variance, our results suggest that the deleterious effects of spatial correlations can be mitigated significantly.

### 4.1. Improved GPCA and TPCA Performance Over MP-PCA

As expected, when spatial correlations are negligible, the novel denoising procedures described here have identical performance to MP-PCA denoising (e.g., Fig 1 and Fig. 3A-B). However, when spatial correlations are introduced (Fig 2 and Fig. 3C-D), our simulations and experiments clearly showed that GPCA and TPCA outperform MP-PCA. The reason for the compromised MP-PCA performance (Fig. 2FG and supplementary Fig. S1) is that 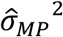 is overestimated by the increased eigenvalue spectral width in the presence of spatially correlated noise; consequently, the criterion for MP-PCA component classification 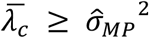 is met only when just a few components are considered as noise carrying. In GPCA on the other hand, since 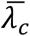 is unaffected by spatial correlations, 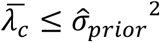 is met as long as 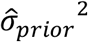 is accurately estimated. For TPCA, 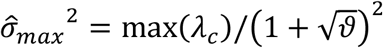 is increased by spatially correlated noise but since the procedure compares 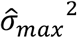 to 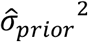, the criterion 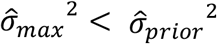 is still satisfied and the noise carrying components are generally correctly labeled.

The limitations of MP-PCA denoising are very clearly noticeable in the pre-clinical data (Fig. 4B), where the introduction of spatially correlated noise is coupled to a very accurate estimation of the noise variance. Therefore, both GPCA and TPCA successfully overcame the MP-PCA limitations. In the human data, MP-PCA denoising also performed sub-optimally (Fig. 6) as highlighted by the deviations between sampled residual quantiles and the theoretical Gaussian quantiles (Fig. 7G). However, since here the noise variance is more difficult to estimate, TPCA provided a better compromise between noise removal and signal preservation. This was observed both directly in diffusion-weighted images (Fig. 4 and Fig. 6), in parametric DKI maps (Fig. 5 and Fig. 7) and in enhanced quantification of high angular information from the high b-value dMRI data (Fig. 8). Taken together, these results illustrate the general benefits of GPCA and TPCA in suppressing noise for advanced diffusion MRI techniques. We expect that these findings can be generalized to other advanced dMRI signal representations, microstructural models, ODF and tractography reconstructions.

### 4.2. Comparison between GPCA and TPCA denoising

Theoretically, GPCA provides a more general eigenvalue classification criteria than MP-PCA and TPCA, since 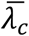 is still a good proxy to the noise variance when components containing mostly noise are properly classified regardless on the specific assumption of the MP distributions (c.f. supplementary material Appendix A and Fig. 2), and thus, one may expect that GPCA would always produce more robust results. This clearly depends strongly on the quality of estimating 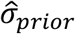. Indeed, when 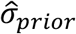 estimates are unbiased, our simulations show that GPCA is the only technique that exactly classifies the correct number of signal and noise components (Fig. 2 and Fig. 3).

However, our work also demonstrates that the performance of GPCA is more sensitive to 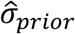 overestimations than the TPCA approach (Fig. 1I and Fig. 2I). This can be explained by the different degrees that the inclusion of eigenvalues related to relevant signal contribution affects the different quantities used by GPCA and TPCA. 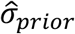 overestimations induce a larger bias in 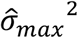 (used by TPCA) than in 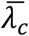 (used by GPCA) as shown in Figs. 1F-G and Figs. 2F-G.

Consequently, TPCA will be more robust to 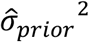 misestimation. This observation is relevant for real life experiments (specially for *in vivo* acquisitions in clinical scanners), since 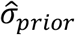 estimation is more likely to be compromised by image artefacts. For instance, in our human diffusion MRI data (Fig. 6A) 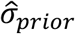 is very likely overestimated, and thus, when GPCA is used, it clearly removed true signal components (Fig. 7A and Fig. 8). Hence, GPCA should be used mainly when 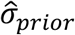 is robustly estimated – e.g. in ex-vivo data where no motion/flow effects exist and where then repetitions can be used more robustly to determine the variance. It is worth noting that the loss of signal due to GPCA denoising can be challenging to observe qualitatively through residual maps (supplementary Fig. S4), which are commonly used for evaluating denoising performance in diffusion MRI. Our results emphasize the need for alternative methods of analysis, such as assessing the impact on diffusion parametric maps that capture non-Gaussian and anisotropic diffusion effects or using residual normality tests to assess denoising performance.

### 4.3. Limitations and Future Work

The “cost” of the new GPCA and TPCA denoising approaches is the need to estimate the noise variance a-priori. In many cases (such as ours) this information can be obtained from just a few repeated b=0 images, which are quite commonly acquired in conventional datasets. Such acquisitions for the common single-shot approaches should cost minimal experimental time and should pose very little issues. This approach avoids strong assumptions on how noise spatially varies across adjacent voxels (Constantinides et al., 1997; Landman et al., 2009b, 2009a; Sodickson et al., 1999). Still, noise variance estimates obtained from repeated images are sensitive to artefacts which can bias 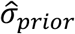 and compromise the correct classification of signal components by our denoising strategies, particularly by GPCA (as discussed above). Moreover, repeated data acquisitions are not always available, especially in retrospective data. Therefore, in future studies, the exploration of alternative noise prior estimation strategies (e.g. (Aja-Fernández et al., 2015, 2009; Landman et al., 2009b, 2009a; Liu et al., 2014; Pieciak et al., 2017; Tabelow et al., 2015)) could be of interest to promote the general use of GPCA and TPCA procedures in different multidimensional MRI datasets.

In this study, our novel denoising strategies are only compared to the original MP-PCA algorithm (Veraart et al., 2016c). Several recent generalization of the MP-PCA denoising have been proposed, as the expansion of MP-PCA to denoise complex data (Bazin et al., 2019; Moeller et al., 2021; Vizioli et al., 2021), higher dimensional MRI data (Olesen et al., 2023), or its integration in multi-coil image reconstruction procedures (Lemberskiy et al., 2021). These expansions are in a sense “orthogonal” to the advances proposed in this study and they can all benefit from the new thresholds given here if a-priori estimates of the noise variance are present. In future studies, GPCA and TPCA component classification can be easily integrated with the above-mentioned (or new) advances in PCA denoising implementations. Additionally, future studies could compare the advantages and disadvantages of our approaches with other denoising strategies that have been emerging in the last years, e.g. algorithms based on machine learning procedures, e.g. (Fadnavis et al., 2020; Muckley et al., 2021).

## 5. Conclusion

The impact of spatially correlated noise on modern reconstructed (vendor) data on the performance of PCA denoising was evaluated and found to corrupt MP-PCA denoising performance. We proposed two new PCA strategies (GPCA and TPCA) that harness prior information on noise variance estimates to objectively denoise MRI data contaminated by spatially correlated noise. Our work shows that diffusion MRI data denoised by both GPCA and TPCA can enhance the quality of diffusion map and orientation distribution function estimates in both pre-clinical and clinical settings. TPCA is more robust from a practical perspective in clinical data since noise variance estimates are likely to be overestimated. Both GPCA and TPCA denoising algorithms are readily generalizable and can therefore be used for denoising other types of redundant data, thereby enabling higher resolution in MRI.

## Acknowledgments

This study was funded by the European Research Council (ERC) (agreement No. 679058). RNH was supported by the Scientific Employment Stimulus 4th Edition from Fundação para a Ciência e Tecnologia, Portugal, ref 2021.02777.CEECIND. AI is supported by “la Caixa” Foundation (ID 100010434) and from the European Union’s Horizon 2020 research and innovation programme under the Marie Skłodowska-Curie grant agreement No 847648, fellowship code CF/BQ/PI20/11760029. The authors thank the University of Minnesota, Magnetic Resonance Research, for the diffusion MRI sequence software used for the brain diffusion data. The authors acknowledge the vivarium of the Champalimaud Centre for the Unknow, a facility of CONGENTO which is a research infrastructure co-financed by Lisboa Regional Operational Programme (Lisboa 2020), under the PORTUGAL 2020 Partnership Agreement through the European Regional Development Fund (ERDF) and Fundação para a ciência e tecnologia (Portugal), project LISBOA-01-0145-FEDER-022170.

