## Supplementary Material for "Efficient PCA denoising of spatially correlated MRI data"

Rafael Neto Henriques[1], Andrada Ianuş[1], Lisa Novello[4], Jorge Jovicich[4], Sune N Jespersen[2,3], Noam Shemesh[1*]

[1]*Champalimaud Research, Champalimaud Foundation, Lisbon, Portugal*
[2]*Center of Functionally Integrative Neuroscience (CFIN) and MINDLab, Clinical Institute, Aarhus University, Aarhus, Denmark.*
[3]*Department of Physics and Astronomy, Aarhus University, Aarhus, Denmark*
[4]*Center for Mind/Brain Sciences - CIMeC, University of Trento, Rovereto, Italy*

*Corresponding author:
Dr. Noam Shemesh, Champalimaud Research, Champalimaud Foundation, Av. Brasilia 1400-038, Lisbon, Portugal

Phone number: +351 210 480 000 ext. #4467

#### Appendix A – General PCA denoising eigenvalue classification

Consider a $M \times N$ matrix $\boldsymbol{X}$ that contains only noise with $\langle X_{ij}\rangle = 0$, $\langle X_{ij}^2\rangle = \sigma^2$, that $\langle X_{ij}X_{kl}\rangle$ can be non-zero even for $(ij) \neq (kl)$, and $M \geq N$, then

$$T = \frac{1}{N}\boldsymbol{X}^T\boldsymbol{X} \Rightarrow$$

$$\langle \frac{1}{N} Tr(T) \rangle = \langle \frac{1}{N}\sum_{i=1}^{N} \lambda_i \rangle = \langle \lambda_i \rangle \Rightarrow$$

$$\langle \frac{1}{N^2} Tr(\boldsymbol{X}^T\boldsymbol{X}) \rangle = \langle \frac{1}{N^2}\sum_{i=1}^{N} (\boldsymbol{X}^T\boldsymbol{X})_{ii}) \rangle = \frac{1}{N^2}\sum_{i,k=1}^{N} \langle X_{ik}X_{ik} \rangle = \frac{1}{N^2}\sum_{i,k=1}^{N} \sigma^2 = \sigma^2$$

Therefore, $\langle \lambda_i \rangle = \sigma^2$ when matrix $\boldsymbol{X}$ contains only noise. When signal components are present $\langle \lambda_i \rangle > \sigma^2$, thus, the eigenvalue classification criterion for General PCA denoising consists in finding the larger number of eigenvalues that satisfies the following inequality:

$$\langle \lambda_c \rangle < \sigma^2$$

Note, that in contrast with the Marčenko-Pastur distribution, here, we do not assume that matrix $\boldsymbol{X}$ has entries with a probability distribution of finite fourth order, and that $M$ and $N \rightarrow \infty$.

### Appendix B – Simulation Supplementary Figures

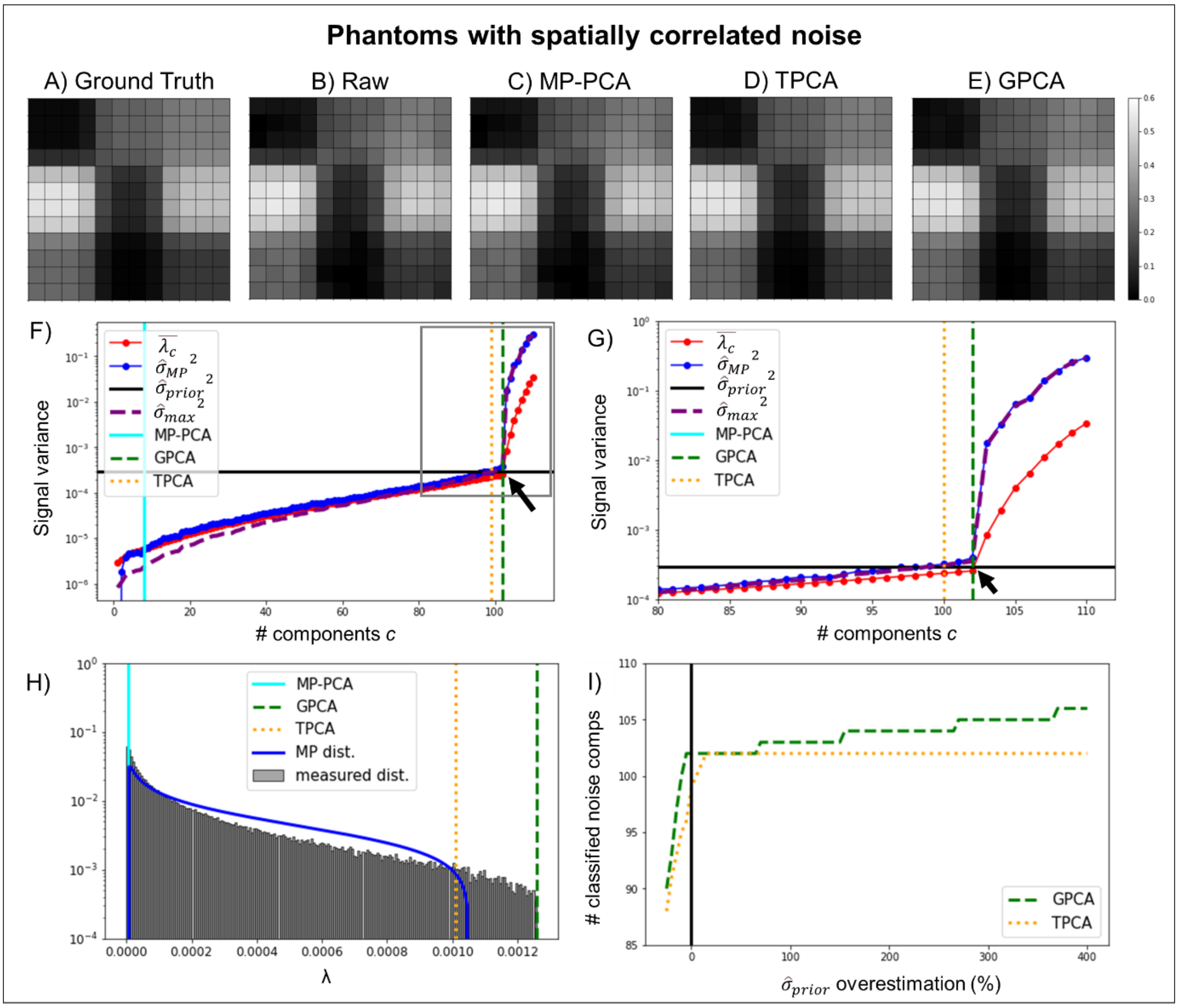


**Supplementary Fig. S1 – Denoising performance in phantom simulation with spatially correlated noise generated by Smoothing data with a 2D Gaussian kernel with a standard deviation of 0.6 (i.e. 34% of signal and noise arising from neigbor voxels).** Representative ground truth noise free and noise corrupted signals for the first diffusion gradient direction of the highest diffusion gradient intensity are shown in panels **(A)** and **(B)** respectively, while denoised signals for the MP-PCA, GPCA and TPCA algorithms are shown in panels **(C)**, **(D)** and **(E)**. Parameters assessed by the denoising algorithms are plotted as a function of the number of lower eigenvalues potentially considered as noise in panel **(F)** - thresholds for the MP-PCA, GPCA and TPCA are plotted by the cyan solid, green dashed, and orange vertical lines respectively (black arrow point to the ground truth number of signal components, i.e., 102). Zoomed plot of the parameters assessed by the denoising algorithms are shown in panel **(G)**. Reconstructed eigenvalue spectrum but running the simulations 1000 times and respective theoretical MP distribution for identical eigenvalue variances are shown in panel **(H)** – the mean thresholds for the MP-PCA, GPCA and TPCA computed as the threshold average across the 1000 repetitions are plotted by the cyan solid, green dashed, and orange vertical lines respectively. The number of classified noise components for both GPCA and TPCA as a function of the percentage overestimation of noise standard deviation is shown in panels **(I)**. In general, this figure shows that the GPCA and TPCA denoising algorithms outperform MP-PCA denoising when noise is spatially correlated.

### Appendix C – Supplementary figures for the pre-clinical data analysis

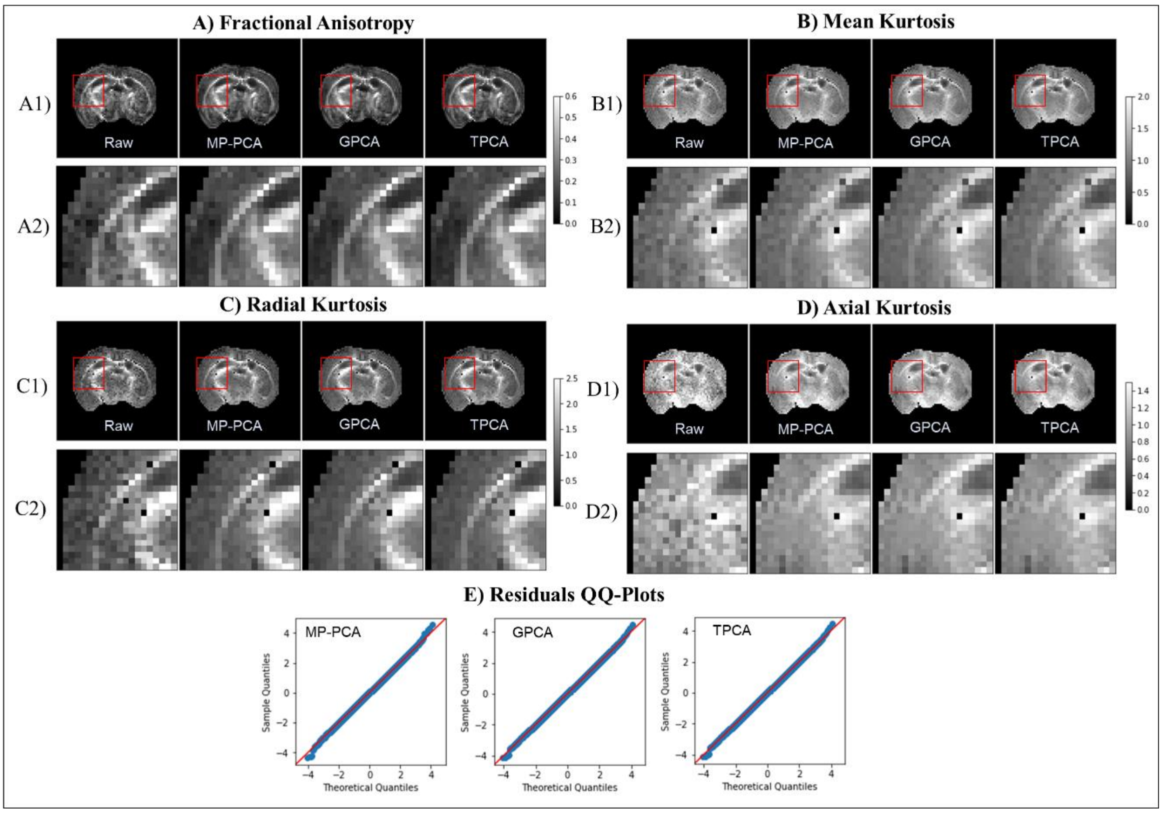


**Supplementary Fig. S2 – DKI maps for the pre-clinical dataset acquired to minimize the effects of spatially correlated noise and QQ-plots of the denoising residuals: A) Fractional Anisotropy; B) Mean Kurtosis; C) Radial Kurtosis; D) Axial Kurtosis; E) QQ-plots of the denoising residuals.** For each DKI quantity, images are displayed for an entire representative axial slice (A1, B1, C1, D1) and for the zoomed area marked by the red box (A2, B2, C2, D2). From left to right, DKI maps are displayed for the raw, MP-PCA denoised, GPCA denoised and TPCA denoised data. QQ-plots of the denoising residuals for the selected zoomed region are show panel E. This supplementary figure shows that, for data acquired with minimized noise spatial correlations, all denoising procedures (MP-PCA, GPCA, TPCA) improves DKI map estimation in a similar fashion. In this case, residual distributions for all denoising techniques is close to a theoretical Gaussian distribution.

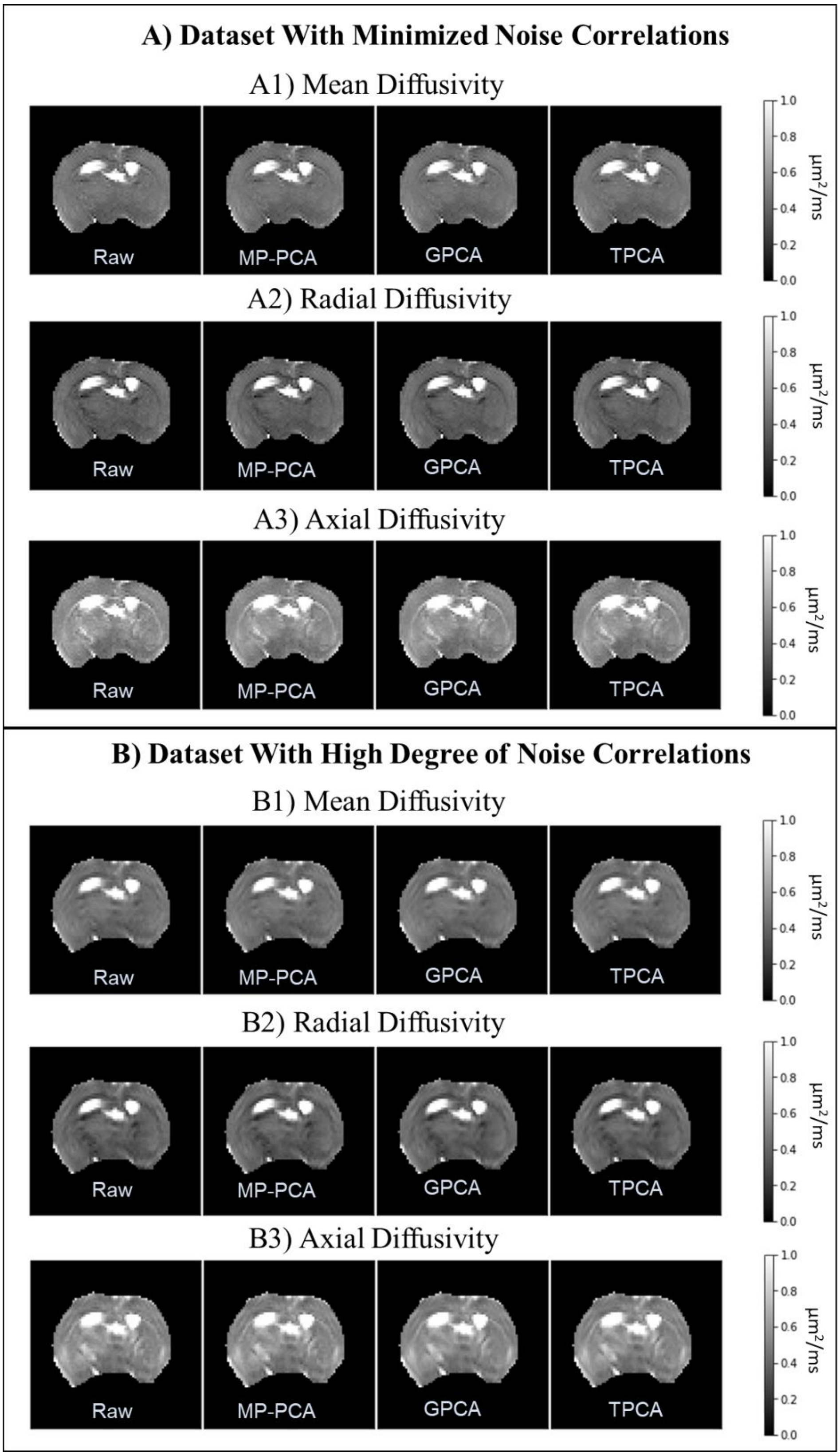


**Supplementary Fig. S3 – DKI diffusivity maps for the two different pre-clinical diffusion MRI datasets: A) dataset acquired with parameters adjusted to minimize noise spatial correlations; and B) typical dataset highly corrupted by spatially correlated noise.** Mean, radial, and axial diffusivities for raw, MP-PCA, GPCA and TPCA are shown in panels (A1/B1), (A2/B2), and (A3/B3) respectively. This figure shows that denoising effects are less evident for diffusion tensor parameters.

### Appendix D – Supplementary figures for the clinical data analysis

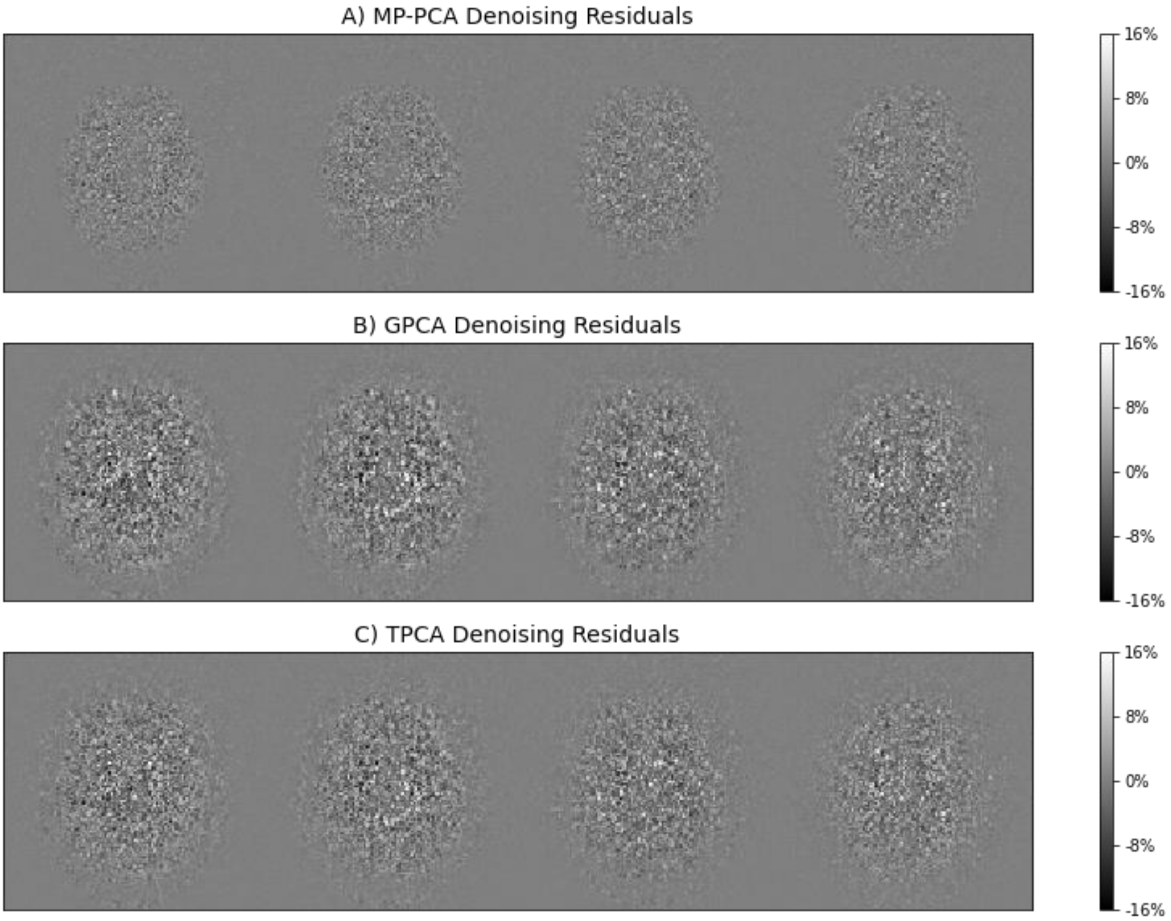


**Supplementary Fig. S4 – Denoising residuals of the clinical dataset for the three different denoising procedures: A)** MP-PCA denoising; **B)** GPCA denoising; and C) TPCA denoising. Residuals are shown for the same representative slices of the main article Figure 6, i.e. residuals are shown for images acquired with gradient direction near v = [1, 0. 0] and for b-value = 2, 3, 4.5, and 6 ms/μm2 (from left to right). This supplementary figure was produced to show that it is hard to inspect loss of structural information on denoising residual maps.
